# Genetic ancestry changes in Stone to Bronze Age transition in the East European plain

**DOI:** 10.1101/2020.07.02.184507

**Authors:** Lehti Saag, Sergey V. Vasilyev, Liivi Varul, Natalia V. Kosorukova, Dmitri V. Gerasimov, Svetlana V. Oshibkina, Samuel J. Griffith, Anu Solnik, Lauri Saag, Eugenia D’Atanasio, Ene Metspalu, Maere Reidla, Siiri Rootsi, Toomas Kivisild, Christiana Lyn Scheib, Kristiina Tambets, Aivar Kriiska, Mait Metspalu

## Abstract

Transition from the Stone to the Bronze Age in Central and Western Europe was a period of major population movements originating from the Ponto-Caspian Steppe. Here, we report new genome-wide sequence data from 28 individuals from the territory north of this source area – from the under-studied Western part of present-day Russia, including Stone Age hunter-gatherers (10,800–4,250 cal BC) and Bronze Age farmers from the Corded Ware complex called Fatyanovo Culture (2,900–2,050 cal BC). We show that Eastern hunter-gatherer ancestry was present in Northwestern Russia already from around 10,000 BC. Furthermore, we see a clear change in ancestry with the arrival of farming – the Fatyanovo Culture individuals were genetically similar to other Corded Ware cultures, carrying a mixture of Steppe and European early farmer ancestry and thus likely originating from a fast migration towards the northeast from somewhere in the vicinity of modern-day Ukraine, which is the closest area where these ancestries coexisted from around 3,000 BC.

## Introduction

The western part of the present territory of Russia has been a focal point of several prehistoric processes yet remains heavily under-represented in ancient DNA (aDNA) studies. Some of the oldest genetically studied individuals from Europe come from this region^1–3^, but overall, ancient genetic information is sparse.

The colonization of the Eastern and Northern European forest belt took place in two large waves during the end of the Paleolithic and the beginning of the Mesolithic period (bordering ca 9,700 cal BC). In both cases, groups of peoples with similar material cultures to those spread in wide areas of Europe took part in the colonization process. In regard to the Mesolithic settlements in the area, a number of distinct archaeological cultures (Butovo, Kunda, Veretye, Suomusjärvi etc.) have been identified^4–7^. In older stages of habitation, the material culture is so similar that it has also been handled as a single cultural area. However, from the middle of the 9th millennium cal BC, local population groups with clearly distinguished cultural differences already existed in the area^8^. Despite a series of small changes occurring during the Mesolithic period (as well as the Early Neolithic period according to the Russian periodization based on pottery production), the cultural continuities as a general trend of those groups are observable across time until the beginning of the 5th millennium, in some areas up to the beginning of the 4th millennium cal BC, when the so-called Pit-Comb Ware and Comb Ware cultures formed in wide areas of Europe^9^. In the territory of the Volga-Oka interfluvial area in Russia, Lyalovo Culture with pit-comb pottery and its local variants were described^10^. It is likely that the people from this cultural realm gave the starting boost to a series of developments in archaeological cultures specific to the 4th to 3rd millenniums cal BC, in particular to the Volosovo Culture, distinguishable in large areas of Russia^11^.

Genetic studies have shown that the Yamnaya Culture people spread out of the Steppe region of the Eastern European Plain and contributed significantly to the ancestry of the European populations^12–14^ that started to produce Corded Ware around 2,900–2,800 cal BC^15^. Furthermore, the migration of the Yamnaya population was two times faster than the Anatolian early farmer (EF) migration into Europe a few thousand years earlier and coincided with a decrease in broad-leaf forests and increase in grasslands/pastures in Western Europe^16^. The Corded Ware Culture (CWC) was spread on a wide area, reaching Tatarstan in the East, the southern parts of Finland, Sweden and Norway in the North, Belgium and Netherlands in the West, and Switzerland and Ukraine in the South^17–19^. Its easternmost extension – Fatyanovo Culture – is a prominent Eastern European CWC branch, that was spread over a large area in European Russia and introduced animal husbandry and probably crop cultivation into the forest belt^20,21^. So far, only 14 radiocarbon dates have been published for Fatyanovo Culture, placing it to 2,750–2,500 (2,300) cal BC^21^. The burial customs characteristic of the culture included the placement of the dead in flat earth graves (less often in barrows), mostly flexed and on their side – men mostly on the right and women on the left side – and shaft-hole stone axes, flint tools and ceramic vessels etc. as grave goods^20,22^.

European Mesolithic hunter-gatherers (HG) can be divided into groups based on their ancestry. The so-called Western group (WHG) was spread from Iberia to the Balkans and reached as far as the Late Mesolithic Eastern Baltic^23–28^. The Eastern group (EHG) had genetic influences from further east (a genetic connection to modern Siberians) and so far includes 6 individuals from Western Russia^14,29,28,30^. The genomes of four of these individuals have been previously studied from Karelia in the northwest from 7,500–5,000 BC^14,28,29^ and two from the Samara region in the eastern part of European Russia from 9,400–5,500 BC^14,30^.

The Yamnaya Culture pastoralists shared ancestry with EHG and Caucasus hunter-gatherers (CHG)^31^. Genetic studies have revealed that CWC individuals, with predominantly Yamnaya ancestry, showed some admixture with European EF of Anatolian ancestry and were most similar to modern populations from Eastern and Northern Europe^13,14,29,32,28^. Lactase persistence, frequent in contemporary Central and Northern Europe, was still at low frequency in the CWC individuals^13,28,29,33^ but underwent strong selection soon after^28,33,34^. It has also been shown that the Yamnaya expansion was male-biased^35^, while the Anatolian EF ancestry in the CWC individuals was acquired more through the female lineage^32^.

In this study, we aim to shed light on the demographic processes accompanying the change from a hunting-gathering lifestyle to crop cultivation and animal husbandry in the forest belt of Northeast Europe and to look into the genetic changes involved in the transition from the Stone to the Bronze Age in the western part of present-day Russia. We add almost 30 new radiocarbon dates from Western Russia and characterize the genetic affinities of the HG and Fatyanovo Culture farmers. As part of the study, we set out to examine if and how the major population movements seen in other parts of Europe during the Holocene have affected this area. Additionally, our aim is to shed light on local processes like the potential admixture between Volosovo and Fatyanovo Culture people suggested by archaeologists^6,17,36^.

## Results

### Samples and archaeological background

In this study, we have extracted DNA from the apical tooth roots of 48 individuals from 18 archaeological sites in modern-day Western Russia and Estonia (Fig. 1, Supplementary Data 1, Supplementary Data 2, Supplementary Note 1). The 28 individuals with better preservation yielded 13– 82% endogenous DNA and <3% contamination (Supplementary Data 2). We have shotgun sequenced these individuals to an average genomic coverage of around 0.1x (n=18), 1x (n=9) and 4x (PES001) (Table 1, Supplementary Data 2).

**Figure 1.**
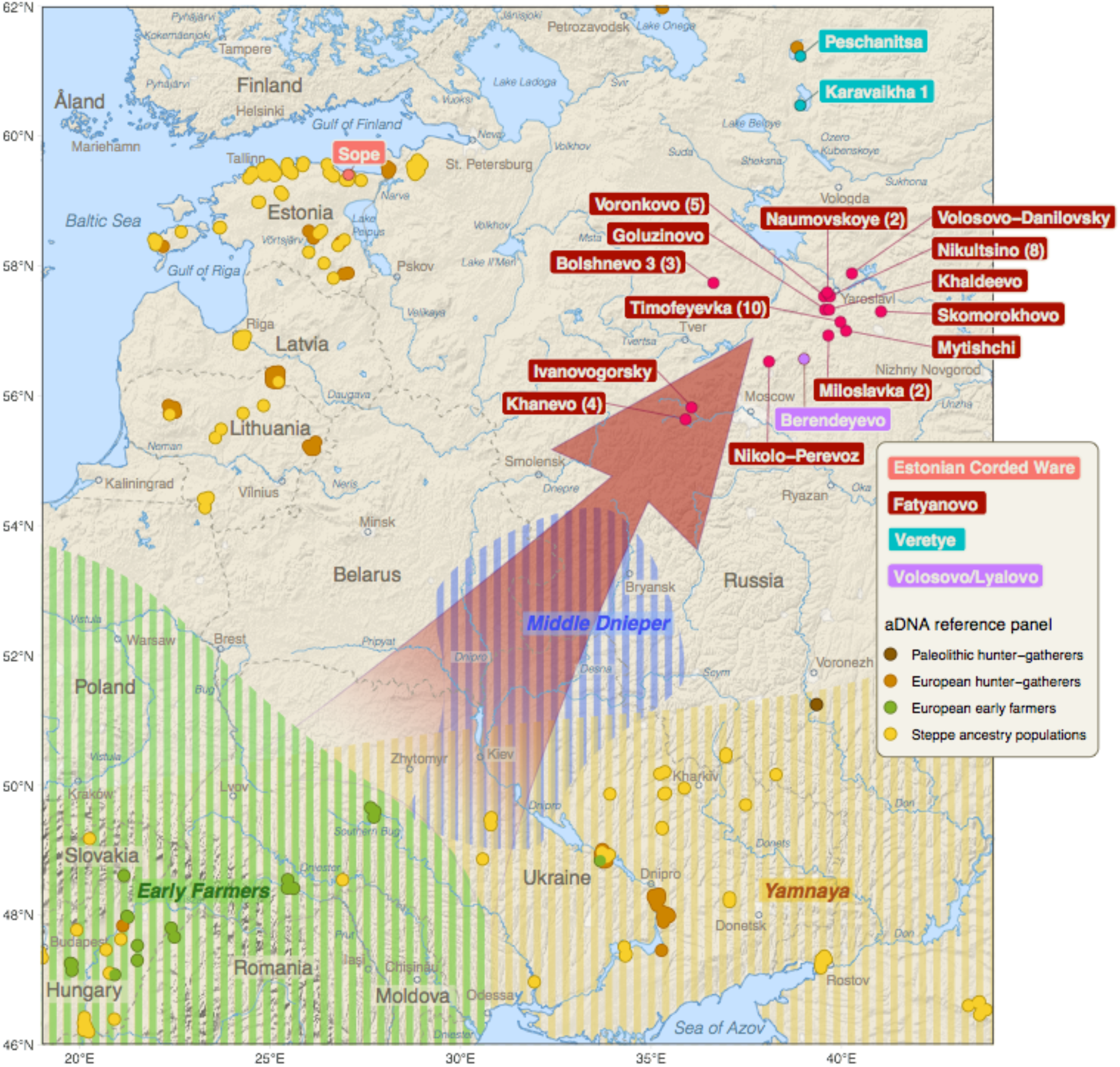
Map of the geographical locations of the individuals of this study. Numbers in brackets behind site names indicate the number of individuals included from this site (if more than one). Arrow indicates the proposed direction of migration of the predecessors of the Fatyanovo Culture people.

**Table 1.**
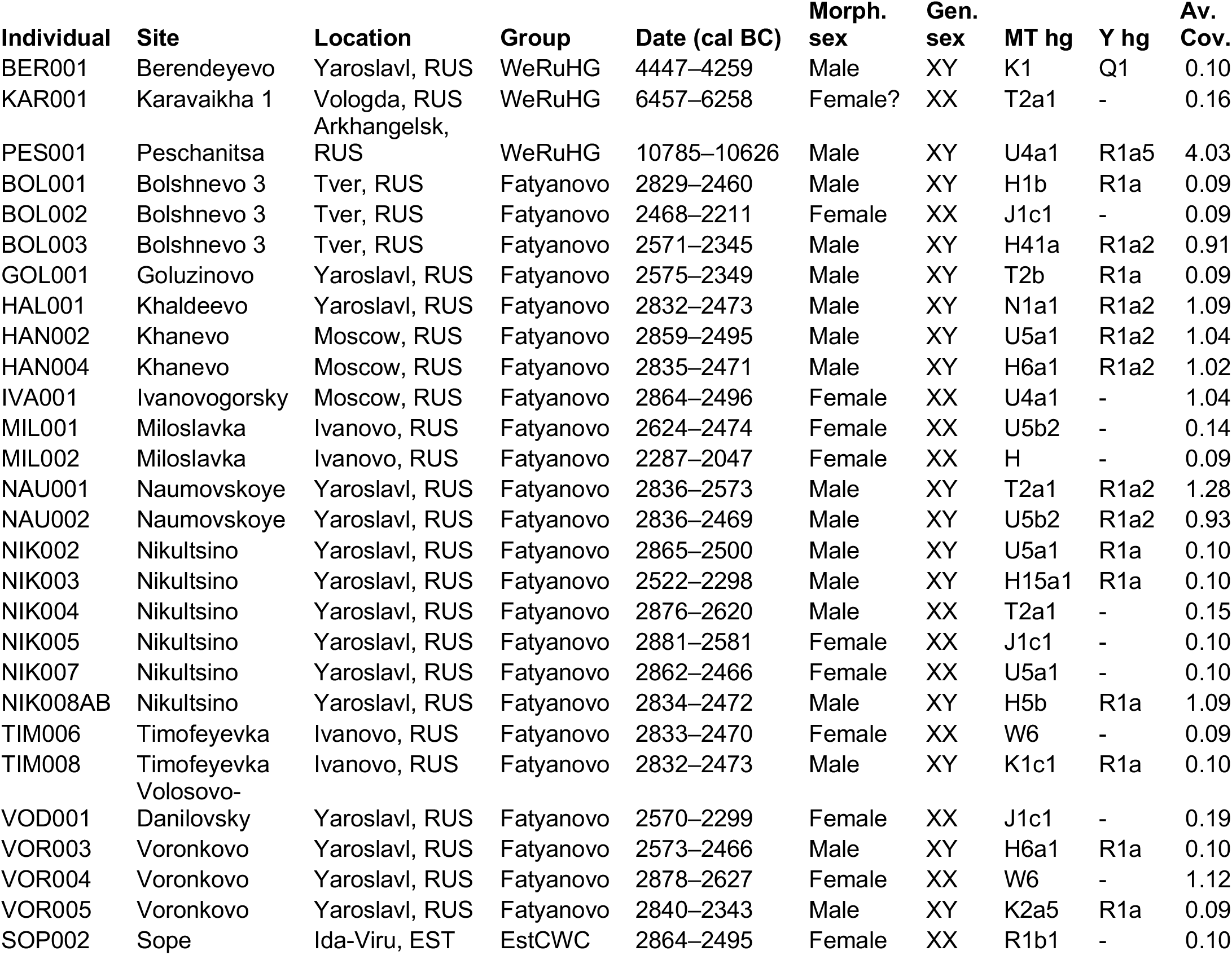
Archaeological information, genetic sex, mtDNA and Y chromosome haplogroups and average genomic coverage of the individuals of this study. Date (cal BC) – calibrated using OxCal v4.2.4^121^ and IntCal13 atmospheric curve^122^; Morph. – morphological; Gen. – genetic; MT hg – mitochondrial DNA haplogroup; Y hg – Y chromosome haplogroup; Av. Cov. – average genomic coverage.

The presented genome-wide data is derived from 3 Stone Age hunter-gatherers (WeRuHG; 10,800–4,250 cal BC) and 24 Bronze Age Fatyanovo Culture farmers from Western Russia (Fatyanovo; 2,900–2,050 cal BC), and 1 Corded Ware Culture individual from Estonia (EstCWC; 2,850–2,500 cal BC) (Fig. 1, Supplementary Data 1, Supplementary Data 2, Supplementary Note 1). We analyzed the data in the context of published ancient and modern populations.

### Affinities of Western Russian hunter-gatherers

First, we assessed the mitochondrial DNA (mtDNA) and Y chromosome (chrY) variation of the 3 Stone Age HG from Western Russia. The oldest individual PES001 belonged to mtDNA haplogroup (hg) U4 (Table 1, Supplementary Data 2), which has been found before in EHG and Scandinavian HG individuals^37,14,27,32,28,26^. The other two represented mtDNA hgs T2 and K1 (Table 1, Supplementary Fig. 5, Supplementary Data 2), which is noteworthy since hg U was by far the most frequent in European HG before the spread of farming, but hgs H11 and T2 have also been found previously in HG individuals ^28,26^. The chrY lineages carried by PES001 and BER001 were R1a5-YP1272 and Q1-L54, respectively (Table 1, Supplementary Data 2) – both hgs have also been found previously in EHG individuals^14,29,32,28,26^.

Next, we compared the WeRuHG individuals to a set of available ancient and modern populations using autosomal data. We performed principal component analysis (PCA), by projecting ancient individuals onto components calculated on Western Eurasian individuals from the Estonian Biocentre Illumina genotyping array dataset (EBC-chipDB). The PCA revealed that all three WeRuHG individuals cluster together with individuals positioned at the EHG end of the European HG cline (Fig. 2A). We then projected ancient individuals onto a world-wide modern sample set from the EBC-chipDB using ADMIXTURE analysis. We ran the calculations on K=3 to K=18 (Supplementary Fig. 1C–D) but discuss K=9 (Fig. 2B, Supplementary Fig. 1A–B). This K level had the largest number of inferred genetic clusters for which >10% of the runs that reached the highest Log Likelihood values yielded very similar results. The analysis again shows that WeRuHG individuals are most similar to EHG, being made up of mostly the blue component maximized in WHG and a considerable proportion of the yellow component most frequent in modern Khanty (Fig. 2B, Supplementary Fig. 1AB).

**Figure 2.**
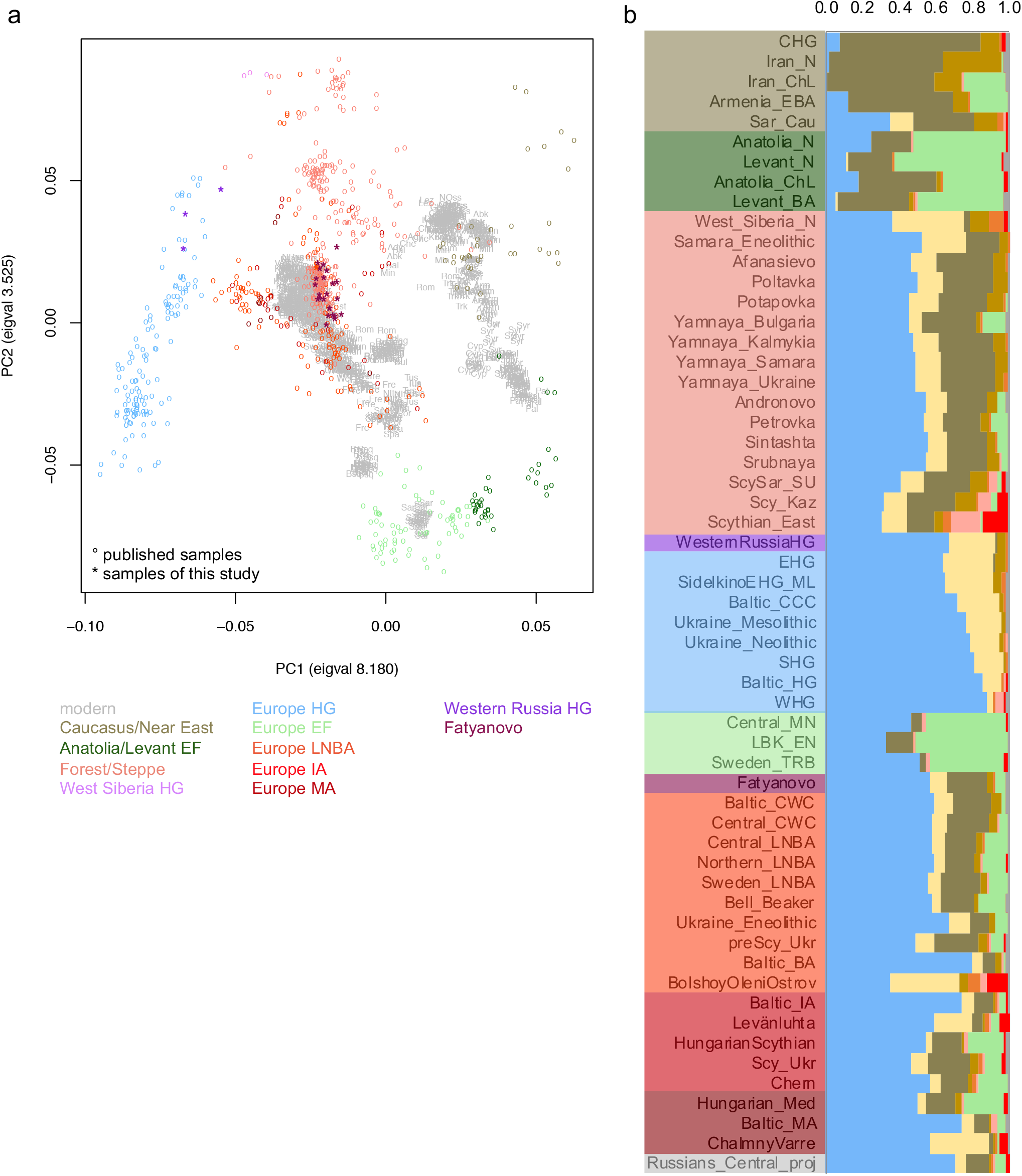
Principal component and ADMIXTURE analyses’ results. **A** principal component analysis results of modern West Eurasians with ancient individuals projected onto the first two components (PC1 and PC2), **B** ADMIXTURE analysis results for a selection of ancient population averages at K9 with ancient individuals projected onto the modern genetic structure. EF – early farmers; HG – hunter-gatherers; LNBA – Late Neolithic/Bronze Age; IA – Iron Age; MA – Middle Ages.

Next, we used outgroup f3 and D statistics to compare the genetic affinity of WeRuHG to those of other relevant populations. We found that WeRuHG and EHG are similar in their genetic affinities both to other ancient and to modern populations (Fig. 3A, Supplementary Fig. 2A). On the other hand, when comparing WeRuHG to the later Fatyanovo, we found that WeRuHG shares more with EHG-like populations, Western Siberian HG, ancient Iranians and modern populations from East Asia and Siberia, while Fatyanovo shares more with most ancient European and Steppe populations and modern populations from the Near East, the Caucasus and Europe (Fig. 3B, Supplementary Fig. 2B).

**Figure 3.**
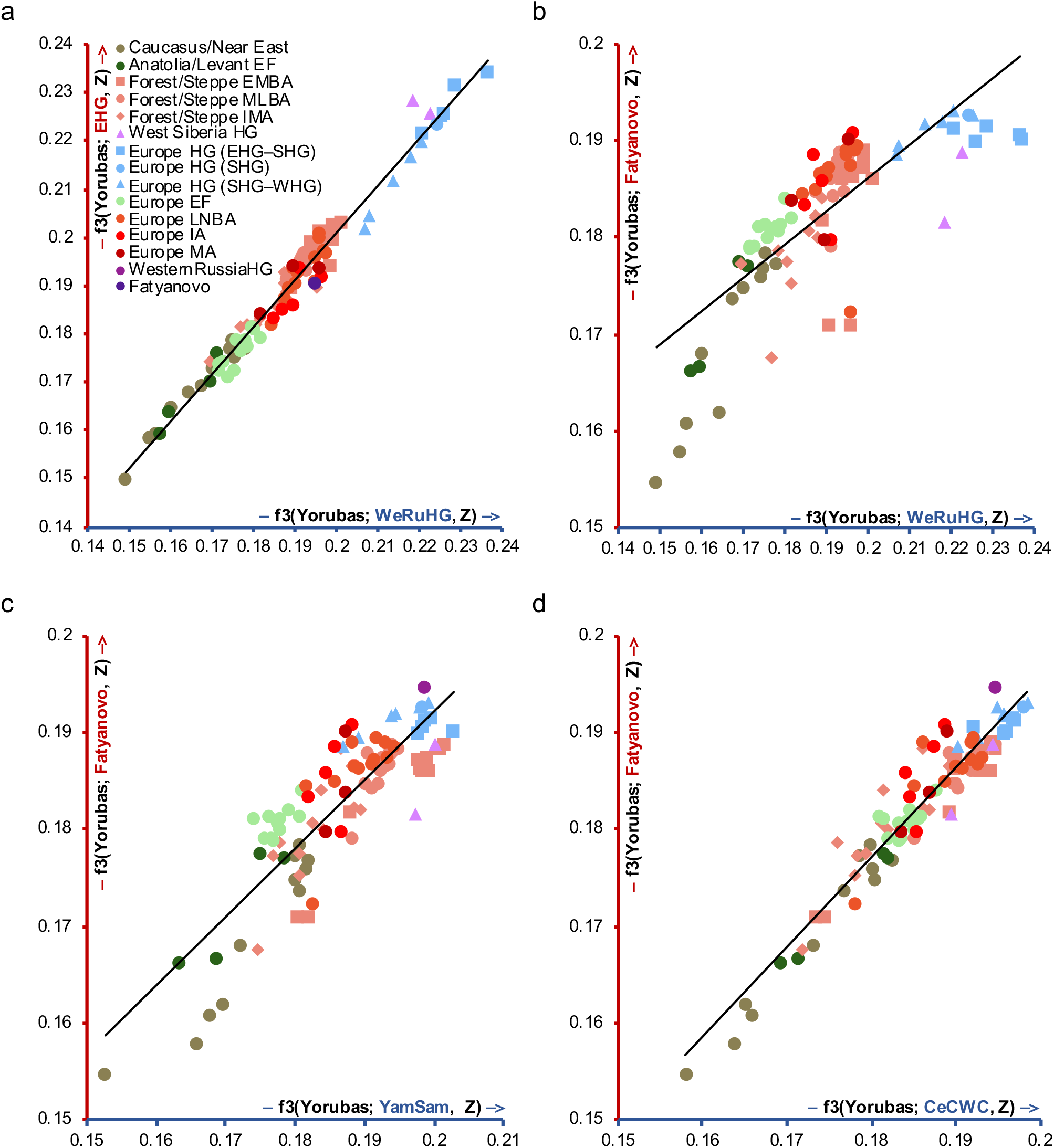
Outgroup f3 statistics’ results of comparisons with ancient populations. Outgroup f3 statistics’ values of form f3(Yorubas; study population, ancient) plotted against each other for two study populations (blue and red axis): **A** Western Russian hunter-gatherers (WeRuHG) and Eastern hunter-gatherers (EHG), **B** WeRuHG and Fatyanovo, **C** Yamnaya Samara (YamSam) and Fatyanovo, **D** Central Corded Ware Culture (CeCWC) and Fatyanovo. EF – early farmers; EMBA – Early/Middle Bronze Age; MLBA – Middle/Late Bronze Age; IMA – Iron/Middle Ages; HG – hunter-gatherers; LNBA – Late Neolithic/Bronze Age; IA – Iron Age; MA – Middle Ages.

### Early farmer ancestry in Fatyanovo Culture individuals

Then, we turned to the Bronze Age Fatyanovo Culture individuals and determined that their maternal (subclades of mtDNA hg U5, U4, U2e, H, T, W, J, K, I and N1a) and paternal (chrY hg R1a-M417) lineages (Table 1, Supplementary Fig. 5, Supplementary Data 2) were ones characteristic of CWC individuals elsewhere in Europe^13,14,29,32,28^. Interestingly, in all individuals for which the chrY hg could be determined with more depth (n=6), it was R1a2-Z93 (Table 1, Supplementary Data 2), a lineage now spread in Central and South Asia, rather than the R1a1-Z283 lineage that is common in Europe^38,39^.

On the PCA, the Fatyanovo individuals (and the Estonian CWC individual) group together with many European Late Neolithic/Bronze Age (LNBA) and Steppe Middle/Late Bronze Age (MLBA) individuals on top of modern Northern and Eastern Europeans (Fig. 2A). This ancient cluster is shifted towards Anatolian and European EF compared to Steppe Early/Middle Bronze Age (EMBA) populations, including the Yamnaya. The same could be seen in ADMIXTURE analysis where the Fatyanovo individuals are most similar to LNBA Steppe ancestry populations from Central Europe, Scandinavia and the Eastern Baltic (Fig. 2B, Supplementary Fig. 1). These populations are composed of the blue “WHG” and yellow “Khanty” component and two brown components maximized in HG from the Caucasus and Iran, similarly to Yamnaya populations. However, the European LNBA populations (including Fatyanovo) also display a green component most frequent in Anatolian and European EF populations, which is not present in the Yamnaya from Russia.

We compared the affinities of the Fatyanovo individuals to those of other populations using f3 and D statistics and found that Fatyanovo shares more with European EF populations and modern Near Easterners than Yamnaya_Samara does (Fig. 3C, Supplementary Fig. 2C, Supplementary Fig. 3C). Importantly, this signal can also be seen when using either autosomal or X chromosome positions from the Lazaridis et al. 2016^31^ ancient dataset instead of the autosomal positions of the EBC-chipDB (Supplementary Fig. 3A–B). However, when comparing Fatyanovo to Central_CWC, there were no clear differences in their affinities to different ancient or modern population groups (Fig. 3D, Supplementary Fig. 2D). Furthermore, we confirmed the presence of sex-biased admixture previously seen in CWC individuals from Estonia, Poland and Germany^40,41,32,42,43^ in the Fatyanovo (Supplementary Fig. 3D). This is also supported by the presence of mtDNA hg N1a in two Fatyanovo individuals – a hg frequent in Linear Pottery Culture (LBK) early farmers, but not found in Yamnaya individuals^44,13,29^.

Since the previous analyses suggested that the genetic makeup of the Fatyanovo Culture individuals was a result of admixture between migrating Yamnaya individuals and contemporary European populations, we used two complementary methods (qpAdm and ChromoPainter/NNLS) to determine the most suitable proxies for the admixing populations and the mixing proportions. The model with the highest p-value and the lowest standard errors for qpAdm had Yamnaya and Levant Neolithic (N) as sources while the model with the smallest residuals for ChromoPainter/NNLS included also WHG, so we are presenting both models for both analyses (Fig. 4A–B). We ran the analyses for Fatyanovo, Central CWC and Baltic CWC and saw that in the results of the more complex model, some of the Yamnaya ancestry from the simpler model got reassigned as WHG but the proportions of Levant N ancestry remained similar (within 4%). We found that although the results from qpAdm and ChromoPainter/NNLS differed somewhat, both showed more Levant N ancestry in Fatyanovo (~22% and ~17%) than in Central/Baltic CWC (~11% and ~10%). What is more, this result is supported by models where Fatyanovo and Central/Baltic CWC are modeled as a mixture of Levant N and Baltic/Central CWC, respectively (Supplementary Fig. 4A).

**Figure 4.**
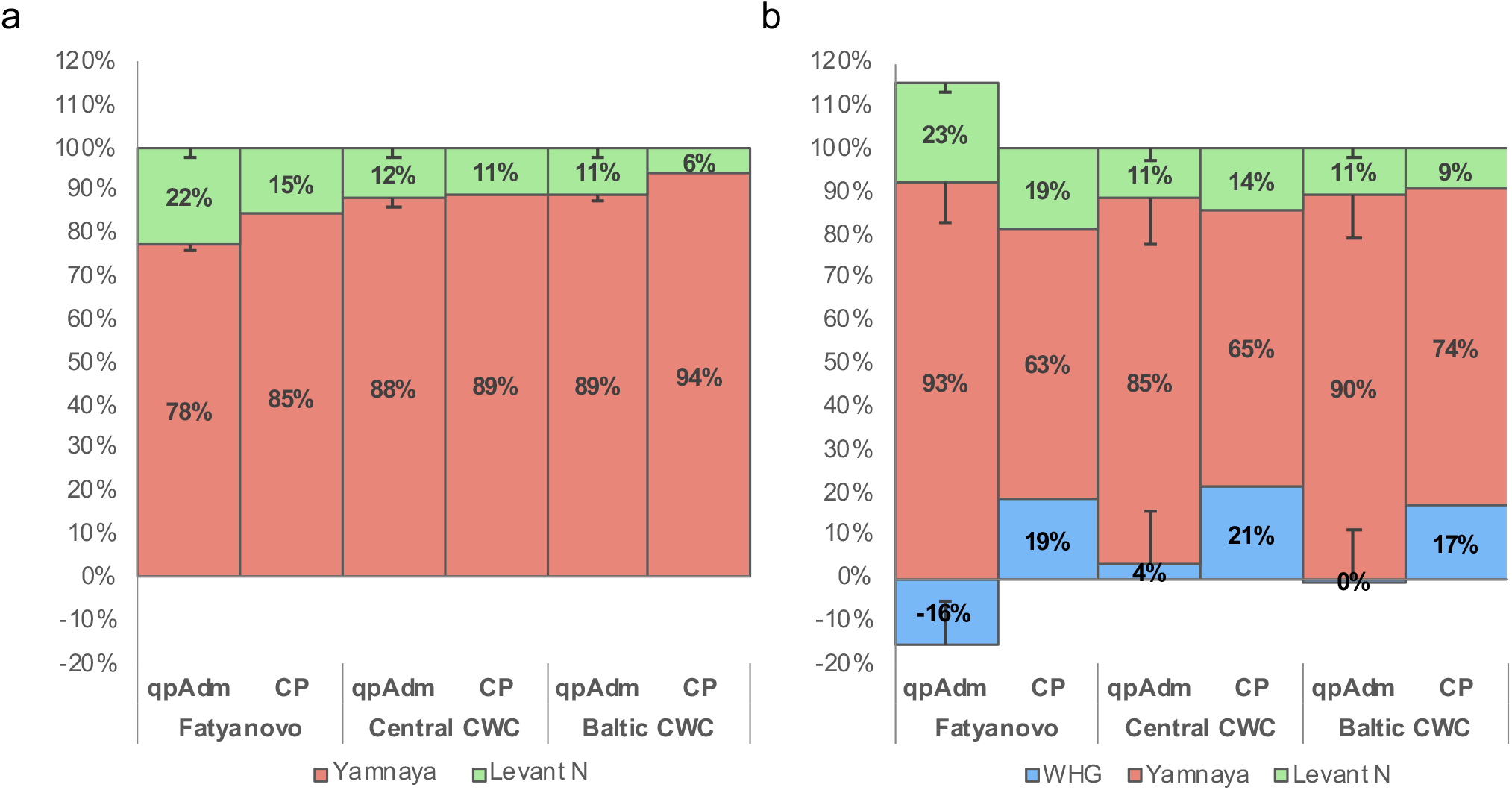
qpAdm and ChromoPainter/NNLS results. **A** models with Yamnaya and Levant Neolithic (Levant N) as sources, **B** models with Western hunter-gatherers (WHG), Yamnaya and Levant N as sources. CP – ChromoPainer/NNLS; CWC – Corded Ware Culture.

Lastly, we looked for closely related individuals in the Fatyanovo Culture sample-set using READ^45^. There were no confirmed cases of 2^nd^ degree or closer relatives (Supplementary Fig. 4B).

### Phenotype informative allele frequency changes in Western Russia

We imputed the genotypes of 115 phenotype-informative positions connected to diet (carbohydrate, lipid and vitamin metabolism), immunity (response to pathogens, autoimmune and other diseases) and pigmentation (eye, hair and skin). We used previously published Eastern Baltic individuals for comparison^27,32,33^. Here, we focus on variants associated with pigmentation (39 SNPs of the HIrisPlex-S system), lactase persistence (rs4988235, rs182549; *MCM6*) and fatty acid metabolism (rs174570C; *FADS2-3*) (Table 2), the latter three alleles showing a significant increase in frequency through time. Although the results should be interpreted with due caution because of the small sample size, we inferred that the examined WeRuHG individuals carried alleles connected to brown eyes, dark brown to black hair and intermediate or dark skin pigmentation while around a third of the Fatyanovo individuals had blue eyes and/or blond hair. Furthermore, we infer that the frequency of the two alleles associated with lactase persistence (rs4988235, rs182549; *MCM6*) is 0% in WeRuHG and ~10% in Fatyanovo Culture individuals (similar to Eastern Baltic populations from the same time periods), but has a significant increase to over 40% by the Late Bronze Age in the Eastern Baltic^33^. On the other hand, an allele connected to an increase in cholesterol and a decrease in triglyceride levels in serum (rs174570C; *FADS2–3*) significantly rises in frequency by the CWC period – from ~20% in Eastern Baltic and Western Russian HG to ~50% in CWC individuals from the two regions. Interestingly, the frequency of the allele increases further in the subsequent time periods in the Eastern Baltic.

**Table 2.** Phenotype prediction results. Phenotype proportions per period.

| Phenotype | Proportion in period |  |  |  |  |  |  |  |
| --- | --- | --- | --- | --- | --- | --- | --- | --- |
|  | Western Russian hunter-gatherers | Latvian hunter-gatherers | Estonian and Latvian Corded Ware Culture farmers | Fatyanovo | Estonian Bronze Age | Estonian Iron Age | Ingrian Iron Age | Estonian Middle Age |
| Blue eyes | 0.00 | 0.80 | 0.29 | 0.25 | 0.70 | 1.00 | 1.00 | 1.00 |
| Brown eyes | 1.00 | 0.20 | 0.71 | 0.75 | 0.30 | 0.00 | 0.00 | 0.00 |
| Blond + Dark blond hair | 0.00 | 0.20 | 0.14 | 0.21 | 0.10 | 0.83 | 1.00 | 0.75 |
| Red hair | 0.00 | 0.00 | 0.00 | 0.00 | 0.00 | 0.00 | 0.00 | 0.25 |
| Brown/Dark brown hair | 0.00 | 0.20 | 0.14 | 0.21 | 0.40 | 0.00 | 0.00 | 0.00 |
| Dark brown + Black hair | 1.00 | 0.60 | 0.71 | 0.58 | 0.50 | 0.17 | 0.00 | 0.00 |
| Very Pale + Pale skin | 0.00 | 0.00 | 0.00 | 0.00 | 0.10 | 0.00 | 0.00 | 0.00 |
| Mixed/Unpredictable Pale-Intermediate skin | 0.00 | 0.00 | 0.00 | 0.08 | 0.10 | 0.00 | 0.67 | 0.50 |
| Intermediate skin | 0.33 | 0.60 | 0.71 | 0.67 | 0.80 | 0.83 | 0.33 | 0.25 |
| Mixed/Unpredictable Intermediate-Dark + Dark + Black skin | 0.67 | 0.40 | 0.43 | 0.25 | 0.00 | 0.17 | 0.00 | 0.25 |
| Lactase persistence | 0.00 | 0.00 | 0.07 | 0.11 | 0.45 | 0.42 | 0.67 | 0.75 |
| Desaturation of omega-3 and omega-6 fatty acids | 0.17 | 0.20 | 0.50 | 0.52 | 0.75 | 0.67 | 1.00 | 0.88 |

## Discussion

After the Last Glacial Period, at the end of the 10th and the beginning of the 9th millennium BC, vast areas in the Eastern Baltic, Finland and northern parts of Russia were populated relatively quickly by hunter-gatherer groups’^46–49^. Flint originating from several places in the Eastern Baltic and the European part of Russia and the similarities in lithic and bone technologies and artefacts suggest the existence of extensive social networks in the forest belt of Eastern and Northern Europe after it was populated^50,51,48^. This has led to the hypothesis that the descendants of Paleolithic HG from Eastern Poland to the central areas of European Russia took part in the process^52,7,53^ and remained connected to their origin, creating a somewhat stretched social network^8^. However, these connections ceased after a few centuries, as can be witnessed from the production of stone tools from mostly local materials, and new geographically smaller social units appeared in the middle of the 9th millennium BC^8^. Ancient DNA studies of this area have included Mesolithic individuals that are from the 8th millennium BC or younger and reveal two genetic groups: WHG in the Eastern Baltic and EHG in Northwestern Russia^14,29,27,32,28,30^. However, no human genomes from the settlement period have been published so far, leaving the genetic ancestry/ancestries of the settlers up for discussion. The individual PES001 from around 10,700 cal BC (probably somewhat affected by the freshwater reservoir effect) presented here with 4x coverage provides evidence for EHG ancestry in Northwestern Russia close to the time it was populated. This, in turn, raises the question of the ancestry of the first people of the Eastern Baltic, keeping in mind the shared social network of the two areas at the time of settlement next to the presence of genetically different groups later in time – a question hopefully answered by future studies.

The formation of Fatyanovo Culture is one of the main factors that affected the population, culture and lifestyle of the previously hunter/fisher-gatherer culture of the Eastern European forest belt. The Fatyanovo Culture people were the first farmers in the area and the arrival of the culture has been associated with migration^17,21^. This is supported by our results as the Stone Age HG and the Bronze Age Fatyanovo individuals are genetically clearly distinguishable. The sample size of our HG is admittedly low, but the three individuals form a genetically homogenous group with previously reported EHG individuals^23–28^ and the newly reported PES001 is the highest coverage whole-genome sequenced EHG individual so far, providing a valuable resource for future studies. What is more, the Fatyanovo Culture individuals (similarly to other CWC people) have mostly Steppe ancestry, but also some EF ancestry which was not present in the area before and thus excludes the northward migration of Yamnaya Culture people with only Steppe ancestry as the source of Fatyanovo Culture population. The strongest connections for Fatyanovo Culture in archaeological material can be seen with the Middle Dnieper Culture^20,54^ spread in present-day Belarus and Ukraine^55,56^. Importantly, the territory of what is now Ukraine is where the most eastern individuals with European EF ancestry and the most western Yamnaya Culture individuals are from based on published genomic data^26,57^ (Fig. 1, Supplementary Data 1). Furthermore, archaeological finds show that LBK reached Western Ukraine around 5,300 BC^58^ and the Yamnaya Culture (burial mounds) arrived in Southeastern Europe around 3,000 BC and spread further as far as Romania, Bulgaria, Serbia and Hungary^59^. These findings suggest present-day Ukraine as the possible origin of the migration leading to the formation of the Fatyanovo Culture and of the Corded Ware cultures in general.

The exact timing of and processes involved in the emergence of the Fatyanovo Culture in European Russia and the local processes following it have also remained unclear. Until recently, the Fatyanovo Culture was thought to have developed later than other CWC groups and over a longer period of time^17,20^. However, radiocarbon dates published last year^21^ and the 25 new dates presented here point towards a fast process, similar in time to CWC people reaching the Eastern Baltic and southern Fennoscandia^60–62^. Importantly, the archaeological cultures are clearly differentiated between the areas. What is more, it has been suggested that the Fatyanovo Culture people admixed with the local Volosovo Culture HG after their arrival in European Russia^6,17,36^. Our results do not support this as they do not reveal more HG ancestry in the Fatyanovo people compared to other CWC groups or any visible change in ancestry proportions during the period covered by our samples (2,900–2,050 BC).

Finally, allele frequency changes in Western Russia and the Eastern Baltic revealed similar patterns in both areas. Interestingly, the frequencies of alleles in the *MCM6* and *FADS2–3* genes, which have been hypothesized to have increased due to dietary changes from the Neolithic onwards^63,29,34^, had significant changes in frequency at different times – the *MCM6* alleles by the CWC period and the *FADS2–3* allele after it. This points toward a more complex scenario for the onset of strong selection on these alleles than just the arrival of farming, as has already been suggested previously^13,29,34,64,28,33^.

## Materials and Methods

### Samples and general information

The teeth used for DNA extraction were obtained with relevant institutional permissions from the Institute of Ethnology and Anthropology of Russian Academy of Sciences (Russia), Cherepovets Museum Association (Russia) and Archaeological Research Collection of Tallinn University (Estonia).

DNA was extracted from the teeth of 48 individuals – 3 from Stone Age hunter-gatherers from Western Russia (WeRuHG; 10,800–4,250 cal BC), 44 from Bronze Age Fatyanovo Culture individuals from Western Russia (Fatyanovo; 2,900–2,050 cal BC) and 1 from a Corded Ware Culture individual from Estonia (EstCWC; 2,850–2,500 cal BC) (Fig. 1, Supplementary Data 1, Supplementary Data 2, Supplementary Note 1). Petrous bones of 13 of the Fatyanovo Culture individuals have been sampled for another project (Kristiansen et al., in preparation). More detailed information about the archaeological periods and the specific sites and burials of this study is given below.

All of the laboratory work was performed in dedicated ancient DNA laboratories of the Institute of Genomics, University of Tartu. The library quantification and sequencing were performed at the Institute of Genomics Core Facility, University of Tartu. The main steps of the laboratory work are detailed below.

### DNA extraction

The teeth of 48 individuals were used to extract DNA. One individual was sampled twice from different teeth.

Apical tooth roots were cut off with a drill and used for extraction since root cementum has been shown to contain more endogenous DNA than crown dentine^65^. The root pieces were used whole to avoid heat damage during powdering with a drill and to reduce the risk of cross-contamination between samples. Contaminants were removed from the surface of tooth roots by soaking in 6% bleach for 5 minutes, then rinsing three times with milli-Q water (Millipore) and lastly soaking in 70% ethanol for 2 minutes, shaking the tubes during each round to dislodge particles. Finally, the samples were left to dry under a UV light for 2 hours.

Next, the samples were weighed, [20 * sample mass (mg)] μl of EDTA and [sample mass (mg) / 2] μl of proteinase K was added and the samples were left to digest for 72 hours on a rotating mixer at 20 °C to compensate for the smaller surface area of the whole root compared to powder. Undigested material was stored for a second DNA extraction if need be.

The DNA solution was concentrated to 250 μl (Vivaspin Turbo 15, 30,000 MWCO PES, Sartorius) and purified in large volume columns (High Pure Viral Nucleic Acid Large Volume Kit, Roche) using 2.5 ml of PB buffer, 1 ml of PE buffer and 100 μl of EB buffer (MinElute PCR Purification Kit, QIAGEN).

### Library preparation

Sequencing libraries were built using NEBNext DNA Library Prep Master Mix Set for 454 (E6070, New England Biolabs) and Illumina-specific adaptors^66^ following established protocols^66–68^. The end repair module was implemented using 30 μl of DNA extract, 12.5 μl of water, 5 μl of buffer and 2.5 μl of enzyme mix, incubating at 20 °C for 30 minutes. The samples were purified using 500 μl PB and 650 μl of PE buffer and eluted in 30 μl EB buffer (MinElute PCR Purification Kit, QIAGEN). The adaptor ligation module was implemented using 10 μl of buffer, 5 μl of T4 ligase and 5 μl of adaptor mix^66^, incubating at 20 °C for 15 minutes. The samples were purified as in the previous step and eluted in 30 μl of EB buffer (MinElute PCR Purification Kit, QIAGEN). The adaptor fill-in module was implemented using 13 μl of water, 5 μl of buffer and 2 μl of Bst DNA polymerase, incubating at 37 °C for 30 and at 80 °C for 20 minutes. The libraries were amplified and both the indexed and universal primer (NEBNext Multiplex Oligos for Illumina, New England Biolabs) were added by PCR using HGS Diamond Taq DNA polymerase (Eurogentec). The samples were purified and eluted in 35 μl of EB buffer (MinElute PCR Purification Kit, QIAGEN). Three verification steps were implemented to make sure library preparation was successful and to measure the concentration of dsDNA/sequencing libraries – fluorometric quantitation (Qubit, Thermo Fisher Scientific), parallel capillary electrophoresis (Fragment Analyser, Agilent Technologies) and qPCR.

One sample (TIM004) had a DNA concentration lower than our threshold for sequencing and was hence excluded, leaving 48 samples from 47 individuals to be sequenced.

### DNA sequencing

DNA was sequenced using the Illumina NextSeq 500 platform with the 75 bp single-end method. Firstly, 15 samples were sequenced together on one flow cell. Later, additional data was generated for some samples to increase coverage.

### Mapping

Before mapping, the sequences of adaptors and indexes and poly-G tales occurring due to the specifics of the NextSeq 500 technology were cut from the ends of DNA sequences using cutadapt 1.11^69^. Sequences shorter than 30 bp were also removed with the same program to avoid random mapping of sequences from other species.

The sequences were mapped to reference sequence GRCh37 (hs37d5) using Burrows-Wheeler Aligner (BWA 0.7.12)^70^ and command mem with re-seeding disabled.

After mapping, the sequences were converted to BAM format and only sequences that mapped to the human genome were kept with samtools 1.3^71^. Next, data from different flow cell lanes was merged and duplicates were removed with picard 2.12 (http://broadinstitute.github.io/picard/index.html). Indels were realigned with GATK 3.5^72^ and lastly, reads with mapping quality under 10 were filtered out with samtools 1.3^71^.

The average endogenous DNA content (proportion of reads mapping to the human genome) for the 48 samples is 31% (Supplementary Data 2). The endogenous DNA content is variable as is common in aDNA studies, ranging from under 1% to over 80% (Supplementary Data 2).

### aDNA authentication

As a result of degrading over time, aDNA can be distinguished from modern DNA by certain characteristics: short fragments and a high frequency of C=>T substitutions at the 5’ ends of sequences due to cytosine deamination. The program mapDamage2.0^73^ was used to estimate the frequency of 5’ C=>T transitions.

mtDNA contamination was estimated using the method from Fu et al. 2013^74^. This included calling an mtDNA consensus sequence based on reads with mapping quality at least 30 and positions with at least 5x coverage, aligning the consensus with 311 other human mtDNA sequences from Fu et al. 2013^74^, mapping the original mtDNA reads to the consensus sequence and running contamMix 1.0-10 with the reads mapping to the consensus and the 312 aligned mtDNA sequences while trimming 7 bases from the ends of reads with the option trimBases.

For the male individuals, contamination was also estimated based on X chromosome using the two contamination estimation methods first described in Rasmussen et al. 2011^75^ and incorporated in the ANGSD software^76^ in the script contamination. R.

The samples show 11% C=>T substitutions at the 5’ ends on average, ranging from 6% to 18% (Supplementary Data 2). The mtDNA contamination point estimate for samples with >5x mtDNA coverage ranges from 0.03% to 1.81% with an average of 0.5% (Supplementary Data 2). The average of the two X chromosome contamination methods of male individuals with average X chromosome coverage >0.1x is between 0.33% and 1.05% with an average of 0.7% (Supplementary Data 2).

### Kinship analysis

A total of 4,375,438 biallelic single nucleotide variant sites, with MAF>0.1 in a set of over 2,000 high coverage genomes of EGC (unpublished; cohort ^77^) were identified and called with ANGSD^76^ command --doHaploCall from the BAM files of the 24 Fatyanovo individuals. The ANGSD output files were converted to .tped format as an input for the analyses with READ script to infer pairs with 1^st^ and 2^nd^ degree relatedness^45^.

The results are reported for the 100 most similar pairs of individuals out of 253 tested and the analysis confirmed that the two samples from one individual (NIK008A, NIK008B) were indeed genetically identical (Supplementary Fig. 3B). The data from the two samples from one individual were merged (NIK008AB) with samtools 1.3^71^ option merge.

### Calculating general statistics and determining genetic sex

Samtools 1.3^71^ option stats was used to determine the number of final reads, average read length, average coverage etc.

Genetic sex was calculated using the script sexing.py from Skoglund et al. 2013^78^, estimating the fraction of reads mapping to Y chromosome out of all reads mapping to either X or Y chromosome.

The average coverage of the whole genome for the samples is between 0.00003x and 4.03x (Supplementary Data 2). Of these, 18 samples have an average coverage of around 0.1x, 9 samples around 1x, one sample around 4x and the rest are lower than 0.02x (Supplementary Data 2). Genetic sexing confirmed morphological sex estimates or provided additional information about the sex of the individuals involved in the study. Genetic sex was estimated for samples with an average genomic coverage >0.005x. The study involves 16 females and 18 males (Table 1, Supplementary Data 2).

### Determining mtDNA haplogroups

The program bcftools^79^ was used to produce VCF files for mitochondrial positions – genotype likelihoods were calculated using the option mpileup and genotype calls were made using the option call. mtDNA haplogroups were determined by submitting the mtDNA VCF files to HaploGrep2^80,81^. Subsequently, the results were checked by looking at all the identified polymorphisms and confirming the haplogroup assignments in PhyloTree^81^.

Haplogroups 41 of the 47 individuals were successfully determined (Table 1, Supplementary Fig. 5, Supplementary Data 2).

### Y chromosome variant calling and haplogroup determination

In total, 113,217 haplogroup informative Y chromosome variants from regions that uniquely map to Y chromosome^38,77,82–84^ were called as haploid from the BAM files of the samples using the --doHaploCall function in ANGSD^76^. Derived and ancestral allele and haplogroup annotations for each of the called variants were added using BEDTools 2.19.0^85^ intersect option. Haplogroup assignments of each individual sample were made by determining the haplogroup with the highest proportion of informative positions called in the derived state in the given sample. Y chromosome haplogrouping was blindly performed on all samples regardless of their sex assignment.

No female samples had reads on the Y chromosome consistent with a haplogroup, indicating that levels of male contamination were negligible. Haplogroups for sixteen (with coverage >0.02x) out of the 18 males were successfully determined (Table 1, Supplementary Data 2).

### Genome-wide variant calling

Genome-wide variants were called with the ANGSD software^76^ command --doHaploCall, sampling a random base for the positions that are present in the EBC-chipDB^86–95^.

### Preparing the datasets for autosomal analyses

The EBC-chipDB^86–95^ was used as the modern DNA background. Individuals from Lazaridis et al. 2016^31^, Jones et al. 2017^27^, Unterländer et al. 2017^96^, Saag et al. 2017^32^, Mittnik et al. 2018^28^, Mathieson et al. 2018^26^, two Damgaard et al. 2018^30,97^ papers, Narasimhan et al. 2018^98^, Lamnidis et al. 2018^99^, Saag et al. 2019^33^ and Järve et al. 2019^57^ were used as the ancient DNA background. The full genome sequencing data of the aDNA background dataset^27,32,97,30,33,57^ in the form of FASTQ files was called as described in the Variant calling section. The 1240k capture data of the aDNA background dataset^26,28,31,96,98,99^ was used in EIGENSTRAT format. The data of the two comparison datasets and of the individuals of this study was converted to BED format using PLINK 1.90 (http://pngu.mgh.harvard.edu/purcell/plink/)^100^, the datasets were merged and 524,920 SNPs of the modern comparison dataset were kept. Individuals with low coverage (<0.02x) were removed from further autosomal analyses, leaving 28 individuals with average genomic coverage >0.09x, resulting in >50,000 SNPs to be used in autosomal analyses. These included 3 from WeRuHG, 24 from Fatyanovo and 1 from EstCWC (Supplementary Data 2).

### Principal component analysis

To prepare for principal component analysis (PCA), a reduced comparison sample-set composed of 817 modern individuals from 46 populations of Europe, Caucasus and Near East and 720 ancient individuals from 110 populations was assembled. The data was converted to EIGENSTRAT format using the program convertf from the EIGENSOFT 7.2.0 package^101^. PCA was performed with the program smartpca from the same package, projecting ancient individuals onto the components constructed based on the modern genotypes using the option lsqproject and trying to account for the shrinkage problem introduced by projecting by using the option autoshrink.

### Admixture analysis

For Admixture analysis^102^, the same ancient sample-set was used as for PCA and the modern sampleset was increased to 1799 individuals from 115 populations from all over the world. The analysis was carried out using ADMIXTURE 1.3^102^ with the P option, projecting ancient individuals into the genetic structure calculated on the modern dataset due to missing data in the ancient samples. The dataset of modern individuals was pruned to decrease linkage disequilibrium using the option indep-pairwise with parameters 1000 250 0.4 in PLINK 1.90 (http://pngu.mgh.harvard.edu/purcell/plink/)^100^. This resulted in a set of 216,398 SNPs. Admixture was run on this set using K=3 to K=18 in 100 replicates. This enabled us to assess convergence of the different models. K=10 and K=9 were the models with the largest number of inferred genetic clusters for which >10% of the runs that reached the highest Log Likelihood values yielded very similar results. This was used as a proxy to assume that the global Likelihood maximum for this particular model was indeed reached. Then the inferred genetic cluster proportions and allele frequencies of the best run at K=9 were used to run Admixture to project the aDNA individuals, for which the intersection with the LD pruned modern dataset yielded data for more than 10,000 SNPs, on the inferred clusters. The same projecting approach was taken for all models for which there is good indication that the global Likelihood maximum was reached (K3–18). We present all ancient individuals on Supplementary Fig. 1 but only population averages on Fig. 2B. The resulting membership proportions to K genetic clusters are sometimes called “ancestry components” which can lead to over-interpretation of the results. The clustering itself is, however, an objective description of genetic structure and as such a valuable tool in population comparisons.

### Outgroup f3 statistics

For calculating autosomal outgroup f3 statistics, the same ancient sample-set as for previous analyses was used and the modern sample-set included 1490 individuals from 92 populations from Europe, Caucasus, Near East, Siberia, Central Asia and East Asia, and Yorubas as outgroup. The data was converted to EIGENSTRAT format using the program convertf from the EIGENSOFT 5.0.2 package^101^. Outgroup f3 statistics of the form f3(Yorubas; West_Siberia_N/EHG/CentralRussiaHG/Fatyanovo/Yamnaya_Samara/Baltic_CWC/Central_CWC, modern/ancient) were computed using the ADMIXTOOLS 1.1^103^ program qp3Pop.

To allow for X chromosome *versus* autosomes comparison, outgroup f3 statistics using X chromosome SNPs were computed. To be able to use the whole ancient comparison dataset for this analysis, the full genome sequencing data of that dataset and the individuals of this study were called as described in the Variant calling section but using the positions of the Lazaridis et al.^31^ ancient dataset. To allow for the use of the bigger number of positions in the ancient over the modern dataset from Lazaridis et al.^31^, Mbuti from Panel C of the Simons Genome Diversity Project^104^ was used as the outgroup. The outgroup f3 analyses of the form f3(Mbuti; West_Siberia_N/EHG/CentralRussiaHG/Fatyanovo/Yamnaya_Samara/Baltic_CWC/Central_CWC, ancient) were run both using 1,055,186 autosomal SNPs and also 49,711 X chromosome positions available in the Lazaridis et al.^31^ ancient dataset. Since all children inherit half of their autosomal material from their father but only female children inherit their X chromosome from their father then in this comparison X chromosome data gives more information about the female and autosomal data about the male ancestors of a population.

The autosomal outgroup f3 results of the two different SNP sets were compared to each other and to the results based on the X chromosome positions of the Lazaridis et al.^31^ ancient dataset to see whether the SNPs used affect the trends seen.

### D statistics

D statistics of the form D(Yorubas, West_Siberia_N/EHG/CentralRussiaHG/Fatyanovo/Yamnaya_Samara/Baltic_CWC/Central_CWC; Russians_Central, modern/ancient) were calculated on the same EBC-chipDB^86–95^ as outgroup f3 statistics. The ADMIXTOOLS 1.1^103^ package program qpDstat was used.

### qpAdm

The ADMIXTOOLS 1.1^103^ package programs qpWave and qpAdm were used to estimate which populations and in which proportions are suitable proxies of admixture to form the populations of this study. Only samples with more than 100,000 SNPs were used in the analyses. The best model tested (taking into account p-values, standard errors and the presence of negative values for proportions) included Fatyanovo/Baltic_CWC/Central_CWC, Yamnaya_Samara and Levant_N as left populations and Han, Yorubas, Chukchis and Ust-Ishim as right populations.

### ChromoPainter/NNLS

In order to infer the admixture proportions of ancient individuals, the ChromoPainter/NNLS pipeline^105–107,33^ was applied. Due to the low coverage of the ancient data, it is not possible to infer haplotypes and the analysis was performed in unlinked mode (option -u). Only samples with more than 20,000 SNPs were used in the analyses. Since ChromoPainter^108^ does not tolerate missing data, every ancient target individual was iteratively painted together with one representative individual from potential source populations as recipients. All the remaining modern individuals from the sample-set used for Admixture analysis were used as donors. Subsequently, we reconstructed the profile of each target individual as a combination of two or more ancient individuals, using the non-negative least square approach. Let *X_g_* and *Y_p_* be vectors summarising the proportion of DNA that source and target individuals copy from each of the modern donor groups as inferred by ChromoPainter. Y_p_ = β_1_X_1_ + β_2_X_2_ + … + β_z_X_z_ was reconstructed using a slight modification of the nnls function in R^109^ and implemented in GlobeTrotter^110^ under the conditions β_g_ ≥ 0 and ∑β_g_ = 1. In order to evaluate the fitness of the NNLS estimation, we inferred the sum of the squared residual for every tested model^111^. The model with the smallest residual values included Yamnaya (Yamnaya^30^), Levant_N (I0867^31^) and WHG (Loschbour^25^) as sources. The resulting painting profiles, which summarise the fraction of the individual’s DNA inherited by each donor individual, were summed over individuals from the same population.

### Phenotyping

The phenotype prediction was performed only on the samples with an average genomic coverage greater than 0.05x, for a total of 28 subjects.

In order to predict eye, hair and skin colour in the ancient individuals, all the 41 variants from 19 genes in 9 autosomes in the HIrisPlex-S system were selected^112–114^ and the region to be analyzed was selected adding 1 Mb at each side of the SNP, collapsing in the same region the variants separated by less than 5 Mb. A total of 10 regions (two for chromosome 15 and one for each of the remaining autosomes) were obtained, ranging from about 6 Mb to about 1.5 Mb. To analyze the other phenotype informative markers (diet, immunity and diseases), 2 Mb around each variant was selected and the overlapping regions were merged, for a total of 47 regions (45 regions in 17 autosomes and 2 regions on the X chromosome). The local imputation pipeline tested and described in (Hui et al., submitted) was used. Briefly, first the variants were called using ATLAS v0.9.0^115^ (task=call and method=MLE commands) at biallelic SNPs with a minor allele frequency (MAF) ≥ 0.1% in a reference panel composed of more than 2,000 high coverage Estonian genomes (EGC) (Pankratov et al., in review; cohort ^77^). The variants were called separately for each sample and merged in one VCF file per chromosomal region. The merged VCFs were used as input for the first step of our imputation pipeline (genotype likelihood update; -gl command on Beagle 4.1^116^), using the EGC panel as reference. Then, the variants with a genotype probability (GP) less than 0.99 were discarded and the missing genotype was imputed with the -gt command of Beagle 5.0^117^ using the large HRC as reference panel^118^, with the exception of variants rs333 and rs2430561, imputed using the 1000 Genomes as reference panel^119^. Finally, a second GP filter was applied to keep variants with GP ≥ 0.85. Then, the 115 phenotype informative SNPs were extracted, recoded and organized in tables, using VCFtools^120^, PLINK 1.9 (http://pngu.mgh.harvard.edu/purcell/plink/)^100^ and R^109^. The HIrisPlex-S variants were uploaded on the HIrisPlex webtool (https://hirisplex.erasmusmc.nl/) to perform the pigmentation prediction, after tabulating them according to the manual of the tool. Two markers of the HIrisPlex-S set, namely the rs312262906 indel and the rare (MAF=0 in the HRC) rs201326893 SNP, were not analyzed because of the difficulties in the imputation of such variants.

The 28 samples analyzed here were compared with 34 ancient samples from surrounding geographical regions from literature, gathering them in 8 groups according to their region and/or culture: a) 3 Western Russian Stone Age hunter-gatherers (present study); b) 5 Latvian Mesolithic hunter-gatherers^27^; c) 7 Estonian and Latvian Corded Ware Culture farmers (present study, ^27,32^); d) 24 Fatyanovo Culture individuals (present study); e) 10 Estonian Bronze Age individuals^33^; f) 6 Estonian Iron Age individuals^33^; g) 3 Ingrian Iron Age individuals^33^; h) 4 Estonian Middle Age individuals^33^. For each variant, an ANOVA test was performed between the 8 groups, applying a Bonferroni’s correction by the number of tested variants to set the significance threshold. For the significant variant, a Tukey test was performed to identify the significant pairs of groups.

## Acknowledgments

This work was funded by research projects of the Estonian Research Council IUT20-7 (A.K.), PRG243 (M.M., La.S., C.L.S., Le.S., A.S.); of the EU European Regional Development Fund 2014-2020.4.01.16-0030, 2014-2020.4.01.16-0125, 2014-2020.4.01.15-0012; Arheograator Ltd. (L.V., Ai.K.). The authors would like to thank the University of Tartu Development Fund for support to the Collegium for Transdisciplinary Studies in Archaeology, Genetics and Linguistics. We thank Marcel Keller for help with coordinates. Analyses were carried out using the facilities of the High Performance Computing Center of the University of Tartu.

## Author contributions

Le.S., A.K. and M.M. conceived the study. S.V.V., L.V., N.V.K., D.V.G. and A.K. assembled skeletal samples and performed osteological analyses. Le.S., S.J.G., K.T., C.L.S. and A.S. performed aDNA extraction and sequencing. Le.S., La.S., E.D’A., E.M., S.R. and T.K. analyzed data. Le.S., L.V., N.V.K., D.V.G., T.K., K.T., A.K. and M.M. wrote the manuscript with input from remaining authors.

