## Supplementary Information for "Genetic ancestry changes in Stone to Bronze Age transition in the East European plain"

### Supplementary Notes

#### Supplementary Note 1. Archaeological and anthropological background

##### Veretye culture

Veretye culture was defined by Svetlana V. Oshibkina based on excavations of Mesolithic sites at the Lacha Lake in the Arkhangelsk region in the Northern part of European Russia (Oshibkina, 1983; 1989; 2006). Excavations of the reference archaeological site of the culture — Veretye 1 – were conducted by Oshibkina in 1978–1991 (Oshibkina 1997), although Mesolithic contexts in the area were studied already in 1920s30s by Mariya Ya. Foss (Foss, 1941) in the Veretye multilayer site (so-called Nizhneye Veretye) that is situated 80 m from Veretye 1. In the end of the 20th and the beginning of the 21st century other sites with similar archaeological materials were discovered and partly excavated in this territory (Oshibkina, 2006).

The distribution of Veretye culture was generally limited by the area of the Vostochnoye Prionezhye region to the east of Lake Onega. This territory consists of glacial lake kettles that are followed now by lake lowlands. Settlements and campsites were located at the river banks often close to the river-mouths at the big lakes. The rather dynamic palaeoenvironmental history of the area (Kvasov 1975; Devyatova 1982; Kosorukova et al. 2018) provided certain conditions as Stone Age archaeological contexts were covered by peat sediments in many cases. Due to the good preservation of organic materials in the archaeological contexts, the material of Veretye culture is diverse, including bone, antler and even wooden artefacts, remains of dwelling and fishing constructions as well as archaeozoological finds etc.

Beside settlements and campsites two burial sites of Veretye culture were studied. Remains of two people accompanied by several anthropogenic structures was discovered from the Peschanitsa burial ground, and 10 graves were excavated in the Popovo burial ground (Oshibkina 1982; 1994; 2017). The burial rites display some parallels to the Oleniy Ostrov Late Mesolithic cemetery in Lake Onega, but there are certain differences in the composition of grave goods (Oshibkina 2017).

The chronology of Veretye culture (Oshibkina 2004) was originally established based on archaeological typology and results of palaeogeographical studies. A series of conventional radiocarbon dates from samples of organic sediments from cultural, overlapping and underlying layers confirmed the rather long chronology of the culture – from the end of the 9th millennium BC until the second half of the 6th millennium BC. However, radiocarbon dates are still relatively few and their context is not clear. The oldest and youngest radiocarbon dates from charcoal and wood found in Veretye 1 settlements, for which the reservoir effect is excluded, are 9,600±80 (Le-1469) – 9,237–8,766 cal BC with a 95.4% probability – and 7,700±80 (Le-1472) – 6,687-6,422 cal BC with a 95.4% probability (Oshibkina 2006). There are also a number of radiocarbon dates from human bones (Oshibkina 2006), but the potential for a reservoir effect must be considered. Two chronological phases were defined – the early one in the end of the Preboreal until the first half of the Boreal climatic periods, and the late one in the end of the Boreal period until the beginning of Athlanticum.

The flint industry of Veretye culture was mainly based on processing local grey moraine flint of rather high quality. Flakes were mainly used as preforms, although blades made up around 20% of the assemblage. Several blade production technological contexts were defined, including large blades, narrow blades and bladelets for slotted tools. Tools made of blades include arrowheads with and without tang, small points, possible knives and one obvious dagger from Veretye 1. Slate, sandstone and some crystalline rocks were used in the Veretye lithic industry as well. The amount of chopping tools made of both flint and slate is quite significant, over 20%. Sandstone spheroids with holes similar to so-called "mace-heads" were also found, widely presented in Mesolithic contexts in Finland and Karelia.

The assemblage of bone and antler tools was presented by a variety of hunting, fishing and household tools, and inset daggers. According to archaeozoological materials, the subsistence of the Veretye culture population was mainly based on exploring forest resources, at least during its early phase. In the later phase, aquatic resources became more sufficient for the economy (Oshibkina 2006).

The origin and quaintness of Veretye culture were recently discussed in a larger context of the Early Holocene cultural history of the Eastern European forest belt. Nowadays it is generally accepted that during the Early Mesolithic time the huge territory of the Eastern European Forest belt was a common cultural space due to the developed system of interregional networks (Oshibkina 2004; Zhilin 2003; Gerasimov et al. 2010). From the middle and end of the 8th millennium BC local cultural peculiarities became more pronounceable (Oshibkina 2006; Kriiska & Gerasimov 2014; Kriiska et al. 1916), which resulted in the formation of different cultural traditions.

Burial sites included in this article

Peschanitsa (Oshibkina 1994; 2006; 2017) burial site is located c 1 km from the eastern shore of the Lacha Lake in the Arkhangelsk region. The site was excavated by S. V. Oshibkina in 1986–87 and 1990–91. Two graves and several pits surrounding them have been discovered. The pits contained flint adzes, scrapers, bladelets, retouched flakes, remnant cores and some bone artefacts, also animal and bird bones covered in red ochre. In one grave, the remains of ribs and a pelvis were found and a skull found by a local inhabitant during gravel mining can most probably be associated with these. Beside the bones, the grave contained three flint bladelets and separated animal and bird bones. The skull is the object of our study (PES001) and it belongs to a 45–50-year-old male. In the second grave, only leg bones were found. Those are radiocarbon dated to 9,890±120 (GIN-4858), 9,983–8,937 cal BC with a 95.4% probability. The bone collections in the two gaves have been considered to belong to both one or two people. Our dating of the skull (10,728±59 (UBA-41633), 10,785–10,626 with a 95.4% probability) confirms that these are two different individuals. In the case of both dates, however, it must be taken into account that they are affected by the freshwater reservoir effect, the magnitude of which we cannot estimate.

Karavaikha 1 multiperiod archaeological site and burial ground is situated on the right bank of the Eloma (a channel of Modlona) river, some 18 km to the south from the Vozhe Lake in the Vologda region. The site was excavated by Alexander Ya. Bryusov during seven field seasons between 1938 and 1955, during which about 580 $m^2$ were unearthed (Bryusov 1941; 1951; 1961; Utkin, Kostyleva 2001). The whole area presents a huge swamp with small patches of ground rising above water some 1–1.5 m. Karavaikha is located on one such small hillwhile the trail of the cultural layer also runs down the hill and was covered by peat and hyttia layers. Archaeological material presents all periods from the Mesolithic to the Bronze Age – there is no stratigraphic sequence, but certain patterns of spatial distribution of finds of different periods has been mentioned (Bryusov 1961; Kosorukova, Piezonka 2014).

A. Bryusov studied 38 ancient graves at the site and four more sites were defined later based on archive data and analysis of the archaeological collection by Alexander V. Utkin and Elena L. Kostyleva (2001), who also suggested an arranged numeration for the graves. The graves were dug in the 30–40 cm thick peat-silt cultural layer or deeper into the underlying clay layer. Burials differ in depth, location on the hill, composition of grave filling and amount of used ochre, which was considered as evidence of their different age. Burial goods are very rare and could instead belong to the cultural layer in general. Five radiocarbon dates associated with the burials (8,200±50 BP (GIN-7173), 7,351–7,066 cal BC with a 95.4% probability; 6,880±90 BP (GIN-7176), 5,978-5,631 cal BC with a 95.4% probability; 5,890±220 BP (GIN-7091), 5,305–4,344 cal BC with a 95.4% probability; 4,760±100 BP (GIN-7090), 3,794–3,127 cal BC with a 95.4% probability; 4,420±50 BP (GN-7172), 3,332–2,915 cal BC with a 95.4% probability) showed a rather wide time range (Utkin, Kostyleva 2001).

Our study includes grave 12, which was excavated in 1939. The grave pit was 46 cm deep, reaching the surface of the clay layer. The grave contained a burial of an adult (KAR001), laid on their back in an extended position, head directed to W, hands extended along the body, head turned to the left (Bryusov 1951). The length of the well-preserved skeleton was 175 cm. The grave filling contained a lot of ochre. Grave 12 has been considered to belong to the earlier phase of the burial ground (the Neolithic), although no archaeological finds can be associated with the burial (Utkin, Kostyleva 2001). Mesolithic (mainly Late Mesolithic) archaeological contexts are well-presented in the nearest neighborhood of the site (Kosorukova, Piezonka 2014).

**Lyalovo and Volosovo cultures**

The Lyalovo culture was defined on the basis of the Lyalovo settlement site near Moscow that was excavated by Boris S. Zhukov (Zhkov 1925). The territory and chronology of the culture has been adjusted multiple times (overview of the research history e.g. Gurina & Krainov 1996; Zaretskaya & Kostyleva 2011). For now, the archaeological sites belonging to the Lyalovo culture have been found between the rivers Volga, Oka and Kostroma in Russia (Gurina & Krainov 1996). Settlement sites belonging to the Lyalovo culture were located on riverbanks and lake shores. Both houses with sunken floors as well as aboveground were built (Gurina & Krainov 1996). Burials can be found at or close to settlement sites (as described by Kostyleva & Utkin 2010). The graves are without inner constructions and typically contain only one inhumation. The deceased were placed in the grave in a supine position, later in a flexed position, and some of them had been either wrapped or tied before the interment. Ochre is known to be found in the graves. The earlier burials can be characterized by a lack of burial goods and embellishments normally found on clothes, except for a few bone artefacts. During the end of the culture period, amber and slate ornaments can be found amongst the artefacts. Lyalovo culture is characterized by large vessels made of clay tempered with mineral admixtures (with ground shell being used as an admixture during the end of the period). The vessels are mainly decorated with pits and comb impressions (Gurina & Krainov 1996; Lozovskiy et al. 2015). In lithic materials, the items of importance include bifacial flint arrow- and spearheads along with polished axes and adzes that were usually made of flint and less often from other rocks (Gurina & Krainov 1996). Bone and wood artefacts, including tools and fishing gear, can be found from settlement sites located in marshlands (Zhilin et al. 2002). The subsistence of the people of Lyalovo culture was based on fishing, hunting and gathering (Gurina & Krainov 1996).

Based on radiocarbon dating, the Lyalovo culture has been placed between ca. 5,200–3,700 cal BC (Zaretskaya & Kostyleva 2011). In the anthropological sense, the Lyalovo culture population is quite heterogenous (Kostyleva & Utkin 2010). The development of the culture (for different opinions on the matter see Gurina & Krainov 1996; Kostyleva & Utkin 2000; Zhilin et al. 2002) has been linked to both the Upper-Volga culture that preceeded the Lyalovo culture and was based in the middle of the European part of Russia (Yu. N. Urban, V. V. Sidorov, A. V. Engovatova), as well as either a widespread or limited migration of small populations from the Northern parts of Russia and Karelia (D. A. Krainov, A. V. Utkin, E. L. Kostyleva).

The Volosovo culture succeeds the Lyalovo culture and is based on the area from Lake Ilmen to the Kama river basin district. Research into this culture group was started by Vasily A. Gordotsov who led the excavation at the Vosolovo settlement site in the Vladimir region in Russia (Gorodtsov 1926). The distribution and the chronology of the culture has been concreticised multiple times (research history can be obtained from Nikitin 1974; Krainov 1987a). It is probable that the Volosovo culture consists of multiple archaeological cultures of the same nature, with the most distinct of them based in the Upper/Middle Volga and Central/Lower Oka regions (Krainov 1987a). The settlements of this culture are situated at the shores of rivers and lakes, with a relatively large number of dwellings researched thus far. The houses may have sunken floors or be above ground and can be either quadrangular or, in fewer cases, slightly rounded (Krainov 1987a; Nikitin 2002; Sidorov 2002b). It is likely that more than one of such houses was simultaneously in use in most villages. The burial sites (as described by Kostyleva & Utkin 2010) can be found within the settlements or nearby. More than 30 burial sites with approximately 250 burials are known. The graves are dug into the ground and have no notable inner constructions. As with the Lyalovo culture, in the Vosolovo culture the dead were laid in the grave in a supine position, a practice which later evolved into burying the dead in a flexed position. The graves are usually meant for single burials and can form rows. Some of the graves included ochre. In the later phase of the culture, the practice of single burials lost its importance and collective graves, as well as partial interments, can be detected. The dead are often buried in clothes that have been lavishly decorated (including amber ornaments) and the grave goods include flint and bone anthropo- and zoomorphic figurines, flint arrowheads etc. However, it has been noted that in some cases the burials do not contain any ornaments or grave goods at all. Ritual practice areas and hoards compiled of different artefacts have also been found from the burial sites.

The artefacts belonging to the Vosolovo culture are diverse. During the earlier period of the culture, vessels were mainly made of clay tempered with shell admixture, which was later replaced by plant-based admixture. In form, the vessels have a rounded bottom (with flat-bottomed forms found in later stages of the culture) and are decorated mainly with pits and comb impressions (Krainov 1987b). In lithic material, small flint artefacts such as scrapers, knives, arrowheads etc., as well as polished axes and adzes are well known in the Volosovo culture (Krainov 1987b) Different tools and fishing gear (such as harpoons, arrow- and spearheads, fishing hooks etc.) made of bone and wood are known from settlements in marshlands. Anthropo- and zoomorphic figurines and amber ornaments are characteristic to this culture (Kashina 2002; Kostyleva & Utkin 2010). The subsistence of the Volosovo culture people was based on freshwater fishing, hunting and gathering (Krainov 1987b; Macāne et al. 2019).

Based on radiocarbon dating, the Vosolovo culture can be placed between the first half of the 4th millennium cal BC to the first half of the 3rd millennium cal BC (discussion see Macāne et al. 2019). The emergence of the culture has been seen as a gradual development in the area between the rivers Volga and Oka (Krainov 1987b; Nikitin 2011), as well as a result of migrations from the area of the Volga-Kama culture (Khalikov 1969) or from the Easter Baltic region (Zhilin et al. 2002). It has also been suggested that the Vosolovo culture may have developed based on the cultural integration of the populations inhabiting the forest belt and the people migrating from the Valdai upland region (Sidorov & Engovatova 1996; Sidorov 2002a).

Burial site included in this article

Berendeyevo (Nikitin 1976; Kostyleva & Utkin 2010) burial site is situated on the edge of the Berendeyevo bog in the Yaroslavl region. Graves with three individuals were discovered during the excavation settlement site Berendeyevo I. Our study includes burial 1, which was unearthed by A. L. Nikitin in 1965. Burial 1 (BER001) is a 55–60-year-old male, who was placed in the grave on his left side in a flexed position, head directed to NW. The size of the grave was 1.1 × 0.7 m and the body had been wrapped in birchbark. Grave goods were not found, but the peat had preserved some remnants of clothing on a few bones and some rope, which had been used to wrap the body. The birchbark of burial 1 has been sampled and two radiocarbon datings were made from the sample: 7,730±40 BP (GIN-197) – 6,638–6,478 cal BC with a 95.4% probability – and 4,340±40 BP (GIN-197b) – 3,086–2,890 cal BC with a 95.4% probability. Birchbark was also discovered and radiocarbon dated in other graves: grave 2 4,970±95 (Mo-446) – 3,972–3537 cal BC with a 95.4% probability – and grave 3 5,540±50 BP (GIN-3886) – 4,487–4271 cal BC with a 95.4% probability. We have dated the individual from burial 1 – 5,487±40 BP (UBA-41612), 4,447–4,259 cal BC with a 95.4% probability. This result is similar to the dating from grave 3, but freshwater reservoir effect cannot be excluded. Burial 1 has been linked to Volosovo culture based on the flexed body position, the birchbark container and the stratigraphy of the site, but according to the radiocarbon date, the Lyalovo culture cannot be ruled out either.

**Fatyanovo culture**

The antiquities of the Fatyanovo culture are recognized in a wide area of the European part of Russia from the Novgorod oblast to Tatarstan, particularly the Lake Ilmen area to the Vyatka River (e.g. Krainov 1987b; Krenke 2019b). The research history of this culture extends back to about 150 years ago when in 1873 the Fatyanovo burial ground, which also gave its name to the culture, was discovered near the town of Jaroslav, prompting the beginning of excavations on the site the same year (Uvarov 1881). The main features of the culture were defined during the first quarter of the 20th century (Gorodtsov 1915). Research was especially active during the 1950s and 1970s when the topic was handled mainly by Dimitri Krainov in the western and Otto Bader in the eastern part of the culture area. Bader (1950) distinguished a separate Balanovo culture, but the two are still commonly approached as a singular comprehensive culture (e.g. Krainov 1987b; Krenke 2019b). Other subgroups have been recognized in the Fatyanovo culture on the basis of differences in pottery and the shape and material of tools as well as burial methods

(Krainov 1972; 1987b). By now, more than three hundred burial places belonging to the Fatyanovo culture (most of them found by chance during other excavation works) and numerous stray finds (mainly shaft-hole axes) have been found.

Information about the settlements of the Fatyanovo culture people is scarce – for a long time, a number of them were only known from the area of the Balanovo group (Krainov 1987b). Found mostly on riverbanks, the settlement sites of the Balanovo group are usually in areas well protected by nature. Both open and fortified settlements were founded by the Balanovo culture group and the remains of dwellings (incl. timber-framed structures), as well as fortifications, have been excavated (Bader & Khalikov 1987). More extensive investigation into the settlement sites of the western part of Fatyonovo culture, including a systematic survey, began in the 2000s (e.g. Krenke 2014; 2019a).

The Fatyanovo culture people buried their dead mostly in inhumation cemeteries without visible above ground signs. In a constrained area of the Middle Volga region, they were buried in singular or grouped barrows, located on higher moraine or sandi hills and the shores of watercourses and lakes (Bader & Khalikov 1987; Krainov 1987). The number of graves in a burial ground varies between 2 and 125 (Krainov 1987). Some regularities in the burial customs can be detected, described in more detail below (Krainov 1972; 1987b; Bader & Khalikov 1987; Volkova 2010). The dead are usually buried at a depth of 0.1–2.7 meters and the grave shafts can be relatively large (maximum 3 × 5.6 meters), some with wooden, birch bark or root constructions. The bodies have been laid in the grave mostly on their sides, with elbow, hip and knee joints flexed. In general, the men are laid on their right side and the women on their left. It has been suggested that some of the deceased had been wrapped prior to burial. As an exception, some cremation burials can be found. In some cases, charcoal and ochre have been found in graves. The grave goods usually consist of pottery vessels, battle axes, flint work axes, adzes and knives, as well as metal weapons and jewelry and in special cases, animal bones and shells. Differences between sexes and age groups have been detected in the depth of the graves as well as the grave goods. The stone battle axes are usually found with male burials, rarely with children and women, with the axe often placed near the head. Polished flint work axes and adzes as well as flint knives are usually found in the hip region in male burials and near the feet in female burials. Other items, such as flint arrowheads, scrapers, blades and flakes, bone tools (awls, points, adzes etc.) and ornaments made of animal teeth, bone and amber can also be found. The age-at-death of the deceased is usually between 30–50 years of age, in special cases up to 70 years. Women, however, are rarely older than 40 years (Krainov 1987b).

The source of subsistence for the people of the Fatyanovo culture has mostly been thought to be animal husbandry, but also hunting, fishing and gathering, and the possibility of cultivation has not been excluded either (Krainov 1972, 1987b). The bones of domestic animals (pig, goat/sheep, sometimes cattle and horse) and items made from the bones have been found from several burial places (Krainov 1987b). Teeth belonging to horses, cattle and goat/sheep have been found from an extensively excavated RANIS settlement site in the Moscow area (Krenke 2019a) and the bones of domesticated animals have also been recovered from many settlement sites of the Balanovo area (Bader & Khalikov 1987). Both nomadic and sedentary animal husbandry has been assumed. The idea of a nomadic lifestyle has been reasoned from the scarcity of settlement sites (e.g. Tretiakov 1966) while the location of the burial places in the landscape is considered as indirect evidence for agriculture (e.g. Bader 1939; Krainov 1987b). The deforestation on the shores of the Moscow River during the Fatyanovo culture period, visible on pollen diagrams, is presumed to be related to agriculture (Ershova 2014). However, direct evidence of cereal crop farming has yet to be found. Hunting as a source of subsistence can be inferred from the bone items found from burial places and animal bones from the settlement sites of the Balanovo group. The bones of elk, roe deer, reindeer, beaver, bear, marten and fox as well as other types of wild animals and fish have been differentiated (Bader & Khalikov 1987; Krainov 1987b).

In the anthropological sense, the Fatyanovo culture people can be seen as a relatively homogeneous group with the greatest similarities, mostly in cranial features, to other Corded Ware cultures (Denisova 1966; Balueva et al. 1988; Varul et al. 2019). Craniometric studies of skulls from the Upper Volga region reveal that the people of Fatyanovo culture are characterized by dolichocranic skulls (long and narrow neurocranium), medium-height and medium-width face, obvious horizontal profiling and a high protruding noseband. These features are especially clear in the oldest skeletons.

Only 14 radiocarbon dates of the Fatyanovo culture sites have been published until this research project and in many cases, the context has been unclear (Krenke 2019b). Based on reliable radiocarbon results, the Fatyanovo culture has been dated to the period between c. 2,750–2,500 (2,300) cal. BC (Krenke 2019b). While our results completely overlap with the beginning of the period, they do shift the end of the culture to be a bit younger, c. 2,150 cal. BC.

Different hypotheses have been proposed on the topic of the origin of Fatyanovo culture as well as on its connections to the cultures of the East European Plain that preceded and followed it. In the first half of the 20th century, the idea of an autochthon emergence of the culture was assumed but many of the researchers later renounced the opinion in favour of migration (Krainov 1987b and references therein). The Fatyanovo culture has been quite unanimously treated as a part of European Corded Ware and Battle Axe cultures. The formation and distribution of the culture has on different occurrences been thought to be both prolonged (Krainov 1987b) as well as a relatively fast process (Krenke 2019b). Mostly, the formation of this culture has been described as a migration of settlers to the area that had previously been inhabited by the people of Vosolovo culture whose main form of subsistence was foraging (Krainov 1987a). The most impactful has been Krainov's (1987) approach where he localized the source of Fatyanovo culture, as well as to all Corded Ware cultures, to the wider area of the basins of rivers Dnepr, Visla and Oder in modern Poland, Ukraine, Belarus, Russia, Lithuania and Latvia. Krainov assumed that the people of the Corded Ware culture settled on the areas of the later Fatyanovo culture in four major migration waves: 1) from Belarus, Lithuania and Eastern Latvia to the Upper-Volga region, 2) from Belarus and Mid- and Upper Dnepr area of Ukraine to Moscow area, 3) from the southernmost areas of Ukraine to Middle-Volga and 4) from Eastern Latvia and Belarus to the areas of Western Dvina and Ilmen. Krainov also did not exclude the idea of some mixing between the settlers and the local populations.

From the pottery found from the Moscow river basin during the 2000s, matches have been found in Globular Amphorae, the pottery of the Rzucewo culture in the eastern coast of the Baltic Sea and the epi Corded Ware of the Southern Poland and Ukrainian Volyn areas. It has been assumed that the Fatyanovo culture developed rapidly on the basis of western Globular Amphorae and Corded Ware culture peoples and that the area of its formulation also included the modern Moscow area (Krenke 2019a; 2019b).

<u>Burial sites included in this article</u>

Bolshnevo 3 (Krainov 1972; 1975) burial site is on a morainal hillock in the Tver region. It is the only Fatyanovo culture burial place in Bolshnevo that has been excavated – the other two which are known were destroyed by gravel mining. D. A. Krainov excavated the site in 1970. He discovered four graves – two were individual graves, while the other two had four skeletons in each grave. We included individual 1 and 2 from grave 4 and the individual from grave 3. A 40–50-year-old male was buried in grave 3 (BOL003). He was buried on his left side in a flexed position, head directed to NE. Grave goods included a battle axe, flint axe, two ceramic vessels, two flint knives, some bone artifacts and a shell. In grave 4 remains of four adults – two male and two female – and a dog were unearthed. Three skulls had signs of perimortem trauma. Individual 1 (BOL001) was a 45–55-year-old male, who was placed on his left side in a flexed position, head directed to E-NE. He had a perimortem injury. The grave goods included a battle axe and a flint knife, also a bear-tooth pendant and a pig's bone were near the skeleton. Individual 2 (BOL002) from grave 4 was a 20–25-year-old woman, who was placed on her left side in a flexed position, head directed to SW. Signs of perimortem trauma were also present on her skeleton. AMS radiocarbon dating results from grave 4 are considerably different (burial 1 4,005±39 BP (UBA-41613), burial 2 3,876±36 BP (UBA-41614)) and conventional date from charcoal (3,960±130 BP (GIN-5240), Krenke 2019b), suggesting that possibly the freshwater reservoir effect should be taken into account in when interpreting the data.

Goluzinovo (Krainov 1972, Krainov & Gadzyatskaya 1987) burial ground is located on a hillock near Sinyavka river in the Yaroslavl region. Five individual graves were excavated by D. Krainov in 1964–65, and burial 4 was included into the current study. The individual (GOL001) is a 20–30-year-old male, who was buried on his right side in a flexed position, head directed to NW. Several grave goods were recovered: a battle axe, a flint work axe, three bone items, a ceramic vessel and half of a shell.

Ivanovogorsky (Krainov 1963; Krainov 1972) burial ground is situated on the shore of the Ruza river in the Moscow region. The burials of two males and one female were unearthed by K. Ya. Vinogradov in 1928–29. We have included burial 3 to our study. The individual (IVA001) is a 30–40-year-old woman, who was buried in a flexed position on her left side, head directed to NE. Grave goods included two ceramic vessels, a flint arrowhead, two flint knives, a whetstone, several bone items, pendants from teeth and tubular bones, and small shells. Pig bones were recovered near her lower limbs – it has been suggested that these are food offerings.

Khaldeevo (Nikitin 1964; Krainov 1972) burial site is on a small morainal hillock in the Yaroslavl region. A. L. Nikitin and D. A. Krainov excavated there in 1959. They discovered one grave, where an 18–25-year-old male (HAL001) was buried. He was placed in a supine position, head directed to SW. Two ceramic vessels, a bone adze and some animal bones were found near the skeleton.

Khanevo (Krainov 1972; Kirianova 1976; Sidorov & Engovatova 1992) burial ground is situated on a hillock on the shore of the Iskona river in the Moscow region. Three graves were excavated by A. V. Sidorov in 1969 and three graves by D. A. Krainov in 1970. Our study includes burials 4, 5, 6 and a cranium (HAN001) which most likely was recovered from a grave that was disturbed by human activity. Burial 4 (HAN004) included a male skeleton aged 18–20 years, who was buried on his right side in a flexed position, head directed to SW. Grave goods included a battle axe, a flint spearhead, a flint arrowhead, a stone axe, a ceramic vessel, a flint knife and two bone items. Additionally, pig bones were recovered – most likely remnants of a food offering. Burial 5 (HAN003) included two individuals, a man and a woman. Both were buried in a flexed position on their right side, heads directed to SW. Several grave goods were found: a battle axe, a stone axe, a flint knife, a flint arrowhead, two ceramic vessels, two bone awls and shells. Near their lower limbs, pig and coat/sheep bones were found – most likely remains from meat that was interred to the grave. Burial 6 (HAN002) is a male aged 18–20 (25) years, who was placed in a supine position, lower limbs flexed from joints, head directed to SW. Grave goods included two work axes, a flint knife, several bone artifacts and a ceramic vessel.

Miloslavka (Krainov 1961; Gadzyatskaya 1964; Krainov 1972) burial site is located on a hillock on the bank of a small lake in the Ivanovo region. D. A. Krainov excavated the site in 1958 and 1961, revealing two graves. Also, reports of two additional destroyed graves were gathered from locals. Three graves out of four were single graves, and one was a double grave. The current study included burials 1 and 4. Burial 1 (MIL001) is a 30–35-year-old female, who was placed in the grave on her left side in a flexed position, head directed to E. Her grave goods included a ceramic vessel, a flint flake and several bone artifacts. Burial 4 (MIL002) is a woman, aged approximately 25 years, who was placed in the grave on her left side in a flexed position, head directed to SE. She was inhumed with a 4–5-year-old child. Several grave goods were recovered: a flint work axe, a ceramic vessel, flint flakes, pendants made from shells, and also animal and bird bones.

Mytishchi (Bader 1959; Gadzyatskaya 1969; Krainov 1972) burial site is situated on a little hillock in the Ivanovo region. One grave with two individuals was excavated by N. P. Milonova and O. N. Bader in 1928 and D. A. Krainov opened another grave in 1966. Our study included material from burial 3 (MOT001), excavated by Krainov – likely an 18–20-year-old woman, who was placed on her left side in a flexed position, head directed to E. The grave had been partially destroyed. Recovered grave goods included a battle axe, four ceramic vessels, several copper items and a bear-tooth pendant.

Naumovskoye (Krainov 1972; Kirianova 1973; Krainov & Gadzyatskaya 1987) burial ground is situated on a hillock next to the Kurbitsa river in the Yaroslavl region. Two graves were unearthed by N. A. Kirianova in 1965. Both individuals are included in the current study. The cranial region of burial 1 was disturbed during gravel mining. The remains of burial 1 (NAU001) derive from a 50–60-year-old male. The individual in burial 2 (NAU002) is an 18–20-year-old male, who was placed into the grave on his left side in a flexed position, head directed to NE. The burial included a battle axe, a ceramic vessel, two flint knives, a bear-tooth pendant, a bone awl, an adze and a circular item made of copper.

Nikolo-Perevoz (Raushenbah 1960; Krainov 1963) is located on the right bank of the Dubna river in the Moscow region. In 1958 V. M. Raushenbah excavated a settlement site which had cultural layers from different time periods and unearthed a mass grave with nine individuals. The heads of six of them were directed to SW and three to NE, eight were placed into the grave in a flexed and one in a supine position. Five ceramic vessels, two battle axes, five arrowheads and a spear head were found. In three instances, some arrowheads characteristic to Volosovo culture were recovered amongst the human remains. Most likely the mass grave included individuals who were killed by Volosovo people. Four skeletons were sampled for this study (RDT001, RDT002, RDT003, RDT004).

Nikultsino (Krainov 1987; Krainov & Gadzyatskaya 1987) burial site is on a hillock in the Yaroslavl region. 18 graves, some with double burials, were excavated by D. A. Krainov in 1964–65. Our study included burials 3b, 6, 7, 9, 11, 15, 16 and 17. Burial 3b (NIK005) was partially destroyed. The individual is a woman, who was placed in the grave on her left side, head directed to NE. A ceramic vessel and shards from two additional ceramic vessels were found. Burial 6 (NIK007) is an 18–20-year-old woman, who was placed in the grave on her left side in a flexed position, head directed to SE. A stone work axe, a flint flake and a lump, numerous pendants from animal teeth, shells and copper, also pig bones and three ceramic vessels were recovered. Burial 7 (NIK008) is a 25–30-year-old male, who was placed in the grave on his right side in a flexed position, head directed to SW. His grave goods included a battle axe, a flint blade, two ceramic vessels, a bear-tooth pendant and a bone awl. Burial 9 (NIK004) is a 50–60-year-old male, who was placed in the grave on his right side in a flexed position, had directed to SSE. His grave goods included a battle axe, a flint work axe, two ceramic vessels, a sheep bone awl and a flint flake. Burial 11 (NIK006) is a 13–16-year-old male, who was placed in the grave on his right side, head directed to SW. A battle axe, a flint knife, two ceramic vessels, flint flakes and quartz flakes were found. Burial 15 (NIK002) is a 40–50-year-old male, who was placed in the grave on his left side in a flexed position, head directed to NE. His grave goods included a battle axe, a flint work axe, a flint knife, a flint arrowhead, a bone pin, two copper items, a grinding stone, a bear-tooth pendant and three ceramic vessels. Burial 16 (NIK003) is a 30–40-year-old male, who was placed in the grave on his left side in a flexed position, head directed to NE. A battle axe was found in the grave. The human remains in burial 17 (NIK001) were poorly preserved and thus age-at-death and sex on the individual have not been assessed. The individual was placed on the left side, most likely in a flexed position. An unidentified flint item, a bone awl, a blank for an awl, a ceramic vessel, animal bones, a piece of a reindeer antler and pieces of red quartzite were found.

Skomorokhovo (Krainov 1972; Gadzyatskaya 1976b) is situated on a sandy hillock on the bank of a small stream which flows to the Zmeyka river in the Ivanovo region. D. A. Krainov excavated three graves and discovered two destroyed graves in 1961–62. Our study includes burial 3 (SKO001) – a 20–30-year-old woman, who was placed in the grave on her right side, head directed to SW. A piece of copper wire and three dog's teeth were found in the grave.

Timofeyevka (Krainov 1964; Krainov 1965; Krainov 1972) burial site is located on a small hillock in the Ivanovo region. In 1959 and 1961, D. A. Krainov excavated 15 single burials and two dog graves. The current study included burials 1, 2, 4, 5, 6, 7, 10, 12, 13, 14 and 15. Burial 1 (TIM007) is a 30–35-year-old male, who was placed on his right side in a flexed position, head directed to SW. The grave goods included a battle axe, a flint knife and a scraper, a ceramic vessel with sheep bones, two flint flakes and some bone artifacts. Burial 2 (TIM006) is a woman with 20–25 (35–40) years of age, who was placed on her left side in a flexed position, head directed to E. Two ceramic vessels, a flint knife, two bone items and sheep bones were found with the individual. Burial 4 (TIM003) is a 35(45)–55-year-old male who was placed in supine position, head directed to SW. He was buried with a flint blade and a flint lump. Burial 5 (TIM004) is a male with 30–40 (45–50) years of age who was placed in a flexed position, head directed to SW. The grave goods included a battle axe, a flint work axe, a flint knife, a fragment of a flint point, a flint scraper, three ceramic vessels, a grinding stone and some bone items. Additionally, five pebbles were placed into the grave. Burial 6 (TIM009) is a (16)18–20-year-old male, who was placed on his right side in a flexed position, head directed to SW. Grave goods included a battle axe, a flint work axe, a boar tusk pendant, a flint point, some bone artifacts, sheep bones and a ceramic vessel. Burial 7 (TIM010) is an 18–20-year-old male, who was placed in the grave on his right side, head directed to SW. A battle axe, flint flakes, a bone dagger, bones from a domestic animal, a round stone and two ceramic vessels were recovered from the grave. Burial 10 (TIM011) is a 12–13-year-old female, who was placed on her left side in a flexed position, head directed to NE. The grave goods were a copper tube, a flint flake, a bone awl, several animal bones and teeth, and a ceramic vessel. Burial 12 (TIM002) is a 12–13-year-old male, who was placed in the grave on his right side in a flexed position, head directed to NW. Two flint flakes, a bone awl and a ceramic vessel were found. Burial 13 (TIM008) is a male with (30–35) 50–60 years of age, who was placed on his right side in a flexed position, head directed to NE. His grave goods included a battle axe, a flint work axe, a flint knife, a ceramic vessel with sheep bones and a bone awl. Burial 14 (TIM005) is a female with 18–20 (25–30) years of age, who was placed in the grave on her left side in a flexed position, head directed to SE. A flint knife, two shell pendants, dozens bone tubular and other beads, several animal and bird bones, and two ceramic vessels were found. Burial 15 (TIM001) is an 18–25(30)-year-old male, who was placed on his right side in a flexed position, head directed to SW. A battle axe, a stone slab, a flint flake and a ceramic vessel with pig and dog bones were found in the grave.

Volosovo-Danilovsky (Krainov 1972; Krainov & Gadzyatskaya 1987) burial site is located on a morainal hillock on the bank of the Levashevk river in the Yaroslavl region. Stray finds have been collected there since the 1930s, archaeological excavations were held by D. A. Krainov in 1962–64. This is the largest known Fatyanovo culture burial place, where 107 graves with 117 individuals have been discovered. Additionally, a dog's burial was found. Previously, one radiocarbon dating of the site has been published, but it is unknown what was dated. The dating – 3,650±80 BP (Le-1044) – 2,281–1,775 cal BC with a 95.4% probability – is disputable. Krenke (2019b) has expressed doubt and the previously published dating also does not correlate with our AMS radiocarbon dating of 2570–2299 cal BC. One individual from burial 11 was included in the aDNA study: an adult woman who was placed on her left side in a flexed position, head directed to NE. Her grave goods included three ceramic vessels, a copper circular item, a flint knife and a work axe.

Voronkovo (Krainov 1972; Gadzyatskaya 1976; Krainov & Gadzyatskaya 1987) burial site is on a morainal hillock on the bank of the Vondeli river in the Yaroslavl region. O.S. Gadzyatskaya excavated 13 graves from the site in 1965. We have included burials 5, 6, 7, 8 and 13. Burial 5 is a 20–30-year-old male, who was placed to the grave on his right side in a flexed position, head directed to SW. A battle axe, a flint work axe, a flint knife, a flint flake and a ceramic vessel were found from the grave. Burial 6 is a 45–55-year-old male, who was placed on his right side in a flexed position, head directed to SW. A battle axe, a flint work axe, two ceramic vessels, a flint knife and other flint, bone and copper items were in the grave. Burial 7 is an 18–20-year-old male, who was placed on his right side in a flexed position, head directed to SW. His grave goods included a battle axe, a flint work axe, a flint knife, a ceramic vessel, and several bone items and shells. Burial 8 is a 50–60-year-old woman, who was placed on her left side in a flexed position, head directed to SW. A battle axe, a flint work axe, a flint knife, three ceramic vessels, numerous tooth pendants, some bone artifacts and flint flakes were discovered near the remains. Also, bird bones were found from the grave backfill. Burial 13 is a 30–40-year-old male, who was placed on his right side in a flexed position, head directed to W. Only a few grave goods were discovered: a flint work axe, a flint flake and one animal bone.

**Estonian regional group of Corded Ware culture**

Judging mainly by the characteristic pottery and stone axes, the Estonian group of Corded Ware culture prevailed in present-day Estonia, northern and eastern parts of Latvia, southeastern Finland, as well as in the Karelian Isthmus, Ingria and the Pskov region in northwestern Russia (Jaanits et al. 1982; Nordqvist 2016, Kriiska et al. 2017). Geographically, it is a neighboring group of Fatyanovo culture, but their contact zone is still almost unexplored. Solitary Fatyanovo-type axes in Estonian Corded Ware culture context and Estonian-type axes in the area of Fatyanovo culture indicate some but not very close contacts between these groups (Jaanits 1973; Nordqvist & Häkälä 2014; Kriiska et al. 2015). Also, in time, the Estonian group is almost analogous to Fatyanovo culture, continuing perhaps only a little longer, the averaged range of AMS dates is 2800–2000 cal BC (Kriiska et al. 2015).

Research on the Estonian Corded Ware culture group began in the second half of the 19th century (Grewingk 1871) and the first general presentations were compiled in the first quarter of the 20th century (Ebert 1913; Tallgren 1922). Several important burial sites were studied in the 1920s and 30s, above all Sope and Ardu (Aul 1935; Indreko 1935; 1937). Excavations at settlement sites started more widely in the 1950s (Jaanits 1955; 19669). The Estonian group is characterized by a relatively large number of settlements (a little less than a hundred) located in different natural environments (Kriiska 2000; Kriiska *et al.* 2015; Paavel *et al.* 2016). Their burials (more than twenty) are all flat earth graves. The graves are simple pits, less than 1.5 m deep and with varying orientations. The burials are single, sometimes double inhumations, the deceased are in a flexed, rarely in a supine position (Indreko 1935; Varul et al. 2019). The grave goods include battle axes, work axes and adzes of flint and crystalline rock, flint knives as well as different bone tools. The subsistence of the Estonian Corded Ware group was likely based on a mixed economy, combining productive livelihoods with hunting, gathering and fishing (Kriiska 2000; Lõugas et al. 2007; Kriiska 2009). Based on aDNA studies (Saag et al. 2017; Mittnik et al. 2018), Corded Ware culture was brought into this area by newcomers, likely coming from somewhere in eastern or central Europe.

Burial site included in this article

Sope (Jaanits et al. 1982; Kriiska et al. 2007; Varul et al. 2019) is a burial ground situated on a sandy knoll on the bank of the Sope stream in Northeastern Estonia. Several graves were exhumed by locals in the end of the 19th and beginning of the 20th century and the unearthed remains were reburied into an unknown location. The site has been archaeologically studied twice: in 1926 by H. Moora who discovered burial Sope I, and in 1933 by R. Indreko who unearthed burial Sope II. According to reported information the burial site included at least 14 individuals, but only remains of Sope I and II have survived. The current study included a sample from Sope II, an adult woman, who was placed into the grave on her right side, lower limbs flexed from joints, head directed towards NW. The burial included a ceramic vessel, a bone awl, a shell of a freshwater pearl mussel (*Margaritifera margaritifera*) and some small round stones.

Bader 1963 = Бадер О. Н. Балановский могильник. Из истории лесного Поволжья в эпоху бронзы. Москва: Издательство Академии наук СССР.

Bader & Khalikov 1987 = Бадер О.Н., Халиков, А.Х. Балановская культура. – Эпоха бронзы лесной полосы СССР. Археология СССР. Москва: Наука, 76–84.

Balueva et al. 1988 = Балуева Т. С., Веселовская Е. В., Лебединская Г. В., Пестряков А. П. Антропологические типы древнего населения на территории СССР. Москва: Наука.

Bryusov 1941 = Брюсов А. Я. Отчет о раскопках 1938-1939 гг. – Труды Государственного Исторического музея, XII. Москва: Государственный исторический музей, 5–20.

Bryusov 1951 = Брюсов А. Я. Свайное поселение на реке Модлоне и другие стоянки в Чарозерском районе Вологодской области. – Ed. М. Е. Фосс Поселения эпохи неолита и раннего металла на севере европейской части СССР. (Материалы и исследования по археологии СССР, 20). Москва, Ленинград: Издательство Академии Наук СССР, 7–76.

Bryusov 1961 = Брюсов, А. Я. Караваевская стоянка. – Ed. А. Я. Брюсов. Сборник по археологии Вологодской области. Вологда: Вологодское книжное издательство, 72–162.

Devyatova 1982 = Девятова Э. И. Природная среда позднего плейстоцена и ее влияние на расселение человека в Северодвинском бассейне и в Карелии. Петрозаводск: Карелия.

Denisova 1966 = Денисова Р. Я. Новые данные об антропологическом типе населения фатьяновской культуры. – Советская этнография, 4 , 24–37.

Ebert, M. 1913. Die baltischen Provinzen Kurland, Livland, Estland. – Praehistorische Zeitschrift V, 498–559.

Ershova 2014 = Ершова, Е.Г. Природное окружение фатьяновских поселений и могильников в долине Москвы-реки по данным спорово-пыльцевого анализа. – Российская Археология, 2014, 4, 19–23.

Foss 1941 = Фосс М. Е. Стоянка Веретье (Отчёт Северной экспедиции ГИМ о раскопках 1929-1934 гг. торфяника в бассейне р. Лача). – Труды Государственного Исторического музея, XII.

Gadzyatskaya 1964 = О. С. Гадзяцкая. Милославский фатьяновский могильник. – Ed. Бадер О. Н. Памятники каменного и бронзового веков Евразии. Москва: Наука,

Gadzyatskaya 1969 = Гадзяцкая, О. С. Фатьяновские могильники в Костромской и Ивановской областях. – Археологические памятники эпохи бронзы на территории СССР Краткие сообщения о докладах и полевых исследованиях Института археологии, 115. Москва: Издательство Академии наук СССР 70–78.

Gadzyatskaya 1976a = О. С. Гадзяцкая. Воронковский фатьяновский могильник. – Восточная Европа в эпоху камня и бронзы. Кольцов Л.В., Зимина М.П., Гадзяцкая О.С. (eds.). Москва: Наука, 129–142.

Gadzyatskaya 1976b = Гадзяцкая О.С. Памятники фатьяновской культуры. Ивановско-Горьковская группа. Археология СССР. Свод археологических источников. Выпуск B1-21. Москва: Наука.

Gerasimov, D.V., Kriiska, A., Lisitsyn, S.N. 2010. Colonization of the Gulf of Finland (the Baltic Sea) coastal zone in the Stone Age. – III Northern Archaeological Congress. Papers. Ekaterinburg - Khanty-Mansiisk: Ural Branch of Russian Academy of Sciences, 29–54.

Gorodtsov 1915 = Городцов В.А. Культуры бронзовой эпохи въ Средней Россiи. – Отчетъ Императорскаго Россiйскаго Историческаго Музея Имени Императора Александра III за 1914 годъ — Москва: Синодальная Типография, 1915, 121–228.

Gorodtsov 1926 = Городцов В.А. Панфиловская палеометаллическая стоянка. – Материалы по изучению Владимирской губернии. Ed. А. И. Иванова. Трудов Владимирского Государственного Областного Музея, II. Владимир: Призыв, 3–20.

Gurina, N. N. & Krainov D. A. 1996. Льяловская культура. – Ошибкина С. В. (Ed.). Неолит Северной Евразии. Москва: Наука, 173–182.

Jaanits, L. 1973. Über die estnischen Bootäxte vom Karlova-Typus. – Finskt Museum 78 (1971), 46–76.

Grewingk, C. 1871. Zur Kenntniss der in Liv-, Est-, Kurland und einigen nachbargegenden aufgefundenen Steinwerkzeuge heidnischer Vorzeit. Verhandlungen der Gelehrten Estnischen Gesellschaft VII, Dorpat.

Indreko, R. 1935. Sépultures néolithiques en Estonie. Õpetatud Eesti Seltsi Aastaraamat 1933, 202–223.

Indreko, R. 1937. Ein Hockergrab in Ardu, Ksp. Kose. Õpetatud Eesti Seltsi toimetised XXX, 185–200.

Jaanits, L. 1955. Neoliitilised asulad Eesti NSV territooriumil. In: Moora, H., Jaanits, L. (eds.). Muistsed asulad ja linnused. Arheoloogiline kogumik I. Tallinn, 176–201.

Jaanits, L. 1966. Venekirveste kultuuri asulatest Eestis. In: Moora, H., Selirand, J. (eds.): Pronksiajast varase feodalismini. Uurimusi Baltimaade ja naaberalade arheoloogiast. Tallinn, 60–66.

Jaanits, L., Laul, S., Lõugas, V., Tõnisson, E. 1982. Eesti esiajalugu. Tallinn.

Kashina, E. A. 2002. Anthropomorphic Flint Sculpture of the European Russian Late Neolithic Age. – Anthropologica et Praehistorica, 113, 51–60.

Khalikov 1969 = Халиков, А. Х. Древняя история Среднего Поволжья. Москва: Наука.

Kirianova 1973 = Кирьянова Н.А. Наумовский фатьяновский могильник. (Ярославская область). – Бронзовый век на территории СССР. (Краткие сообщения Института археологии, 134), 73–77.

Kirianova 1976 = Кирьянова. Н.А. Фатьяновский могильник у д. Ханево. – Восточная Европа в эпоху камня и бронзы. Кольцов Л.В., Зимина М.П., Гадзяцкая О.С. (eds.). Москва: Наука, 143–154.

Kosorukova & Piezonka 2014 = Косорукова Н. В. Пьецонка Х. Новые материалы по эпохе неолита в бассейне озера Воже. – Археология озерных поселений: хронология культур и природно-климатические ритмы. СПб: Археология озерных поселений: хронология культур и природно-климатические ритмы. Санкт-Петербурга: Периферия, 2014, 69–174.

Kosorukova et al. 2018 = Н. В. Косорукова, М. А. Кулькова, Х. Пиецонка. Хронология раннего неолита бассейна озера Воже. – Уральский исторический вестник, 3 (60), 78–86.

Kostyleva & Utkin 2000 = Костылева Е. Л., Уткин А. В. К вопросу о происхождении Льяловской культуры. – Юбилейный сборник, посвящённый 85-летию со дня рождения профессора Ю. А. Якобсона. Иваново, 19–23.

Kostyleva & Utkin 2010 = Костылёва Е.Л., Уткин А.В. Нео-энеолитические могильники Верхнего Поволжья и Волго-Окского междуречья. Планиграфические и хронологические структуры. Москва: ТАУС.

Krainov 1961 = Крайнов. Д. А. Новые памятники фатьяновской культуры. – Краткие сообщения о докладах и полевых исследованиях Института археологии, 84, 83–90.

Krainov 1963 = Крайнов Д.А. Памятники фатьяновской культуры. Московская группа. Свод археологических источников. Археология СССР. Выпуск В 1-19. Москва: Изд-во Академии наук СССР.

Krainov 1964 = Крайнов, Д. А. Памятники фатьяновской культуры. Ярославско-Калининская группа. Свод археологических источников. Археология СССР. Выпуск В 1-20. Москва: Изд-во Академии наук СССР.

Krainov 1965. = Крайнов Д.А. Новые исследования памятников фатьяновской культуры – Новое в советской археологии. Памяти Сергея Владимировича Киселёва. К 60-летию со дня рождения. (Материалы и исследования по археологии СССР/ Акад. наук СССР. Ин-т археологии,130) Ed. Е.И. Крупнов. Москва: Наука, 115–118.

Krainov 1972 = Крайнов. Д.А. Древнейшая история Волго-Окского междуречья. Москва: Наука.

Krainov 1975 = Крайнов Д.А. Болшневский могильник фатьяновской культуры // Памятники древнейшей истории Евразии / Отв. ред. П.М.Кожин, Л.В. Кольцов, М.П. Зимина. Москва: Наука, 1975. С. 173–180.

Krainov 1987a = Крайнов Д.А. Волосовская культура. – Эпоха бронзы лесной полосы СССР. Археология СССР. Москва: Наука, 10–28.

Krainov 1987b = Крайнов Д.А. Фатьяновская культура. – Эпоха бронзы лесной полосы СССР. Археология СССР. Москва: Наука, 58–76.

Krainov & Gadzyatskaya 1987 = Крайнов Д.А., Гадзяцкая О.С. Фатьяновская культура. Ярославское Поволжье. Свод археологических источников. Археология СССР. Выпуск В 1-22. Москва: Наука.

Kriiska A., 2000. Corded Ware Culture Sites in North-Eastern Estonia. − De temporibus antiquissimus ad honorem Lembit Jaanits. Eds. V. Lang & A. Kriiska. (Muinasaja teadus, 8), Ajaloo Instituut, Tallinn, 59–79.

Kriiska, A. 2009. The Beginning of Farming in the Eastern Baltic Area. In: Dolukhanov, P.M., Sarson, G.R., Shukurov, A.M. (eds.): The East European Plain on the Eve of Agriculture. British Archaeological Reports International Series 1964, Oxford, 159–179.

Kriiska & Gerasimov 2014 = Крийска А., Герасимов Д. В. Период позднего мезолита в восточной части Балтийского моря: формирование берегового расселения от Рижского до Выборгского. Ed. В. Н. Карманов. От Балтики до Урала: изыскания по археологии каменного века). Сыктывкар: Институт языка, литературы и истории Коми научного центра Уральского отделения Российской академии наук, 5−36.

Koltsov L.V, Zhilin M.G. 1999 Tanged points cultures in the upper Volga Basin. – Eds. S. K. Kozłowski & J. Gurba. Tanged Points Cultures in Europe. Read at the International Archaeological Symposium. Lublin, September 13-16, 1993. Lublin: Maria Curie-Skłodowska University Press, 346–360.

Kvasov 1975 = Квасов Д. Д. Позднечетвертичная история круп ных озер и внутренних морей Восточной Европы. Ленинград: Наука.

Krenke 2014 = Кренке Н. А. Москворецкие памятники фатьяновской культуры. – Российская Археология, 4, 5–18.

Krenke 2019a = Кренке Н. А. Древности бассейна Москвы-реки от неолита до Средневековья: этапы культурного развития, формирование производящей экономики и антропогенного ландшафта

Krenke 2019b = Кренке Н. А. Радиоуглеродная хронология фатьяновской культуры. – Российская археология, 2, 110–116.

Kriiska, A., Nordqvist, K., Gerasimov, D.V., Sandell, S. 2015. Preliminary Results of the Research at Corded Ware Sites in the Narva–Luga Interfluve, Estonian–Russian Border Area in 2008–2014. Archaeological Fieldwork in Estonia 2014, 39–50.

Kriiska et al. 2017 = Крийска, А, Нордквист, К, Герасимов Д.В. Эстонский вариант шнуровой керамики. V (XXI) всероссийский археологический съезд. Сборник научных трудов. Eds. А. П. Деревянко, А. А. Тишкин. Барнаул: Алтайский государственный университет, 557−558.

Lozovskiy et al. 2015 = Лозовский В. М., Лозовская О. В., Зайцева Г. И., Климов А. А. Комплекс ямочно-гребенчатой керамики льяловского горизонта стоянки Замостье 2. – Неолитические культуры Восточной Европы: хронология, палеоэкология, традиции. Материалы международной научной конференции, посвященной 75-летию Виктора Петровича Третьякова, Санкт-Петербург, 12-16 мая 2015 г. Санкт-Петербург: Институт истории материальной культуры Российской Академии наук, 97–100.

Lõugas, L., Kriiska, A., Maldre, L. 2007. New Dates for the Late Neolithic Corded Ware Culture Burials and Early Husbandry in the East Baltic Region. Archaeofauna 16, 21−31.

Macāne, A., Nordqvist, K. & Kostyleva, E. 2019. Marmot incisors and bear tooth pendants in Volosovo hunter-gatherer burials. New radiocarbon and stable isotope data from the Sakhtysh complex, Upper-Volga regioon. – Journal of Archaeological Science: Reports, 26. https://doi.org/10.1016/j.jasrep.2019.101908

Mittnik, A., Wang, C.-C., Pfrengle, S., Daubaras, M., Zarina, G., Hallgren, F., Allmäe, R., Khartanovich, V., Moiseyev, V., Tõrv, M., Furtwängler, A., Andrades Valtueña, A., Feldman, M., Economou, C., Oinonen, M., Vasks, A., Balanovska, E., Reich, D., Jankauskas, R., Haak, W., Schiffels, S., Krause, J. 2018. The Genetic Prehistory of the Baltic Sea Region. Nature Communications 442 (9), 1–11.

Nikitin 1964 = Никитин А.Л. Фатьяновский могильник у с. Халдеево. (Ростовский район Ярославской области). – Советская археология, 3, 277–282.

Nikitin 1974 = Никитин, А.Л. Существовала ли «волосовская культура». – Советская археология, 2, 27–33.

Nikitin 1976 = Никитин А.Л. Неолитическое поселение Берендеево I. (Ярославская обл.). – Советская археология, 3, 191–202.

Nikitin 2002 = Никитин, В. В. Эволюция жилищ эпох камня и раннего металла в Марийском Поволжье. – И.Н. Черных (ed.) Тверской археологический сборник, 5. Тверская областная типография, Тверь, 363–369.

Nikitin 2011 = Никитин В. В. Поздний неолит в лесной полосе бассейна Волги (к проблеме истоков волосовской культуры и её локальных вариантов). – И.Н. Черных (ed.) Тверской археологический сборник, 8 (1). Тверь: ООО «Издательство «Триада», 213–219.

Nordqvist, K. 2016. From Separation to Interaction: Corded Ware in the eastern Gulf of Finland. Acta Archaeologica 87 (1), 49–84.

Nordqvist, K., Häkälä, P. 2014. Distribution of Corded Ware in the Areas North of the Gulf of Finland: An update. Estonian Journal of Archaeology 18 (1), 3–29.

Sidorov 2002a = Сидоров В. В. Волосовская культура во II тысячилетии до н.э. – И.Н. Черных (ed.) Тверской археологический сборник, 5. Тверская областная типография, Тверь, 363–369.

Sidorov 2002b = Сидоров В. В. Волосовские жилища поселений Маслова Болота. – И.Н. Черных (ed.) Тверской археологический сборник, 5. Тверская областная типография, Тверь, 322-331.

Sidorov & Engovatova 1996 = Сидоров В.В., Энговатова А.В. 1996. Протоволосовский этап или культура? – И.Н. Черных (ed.) Тверской археологический сборник, 2. Тверь, 164–181.

Oshibkina 1982 = Ошибкина С. В. Мезолитический могильник «Попово» на р. Кинеме. – Советская археология, 3, 122-138.

Oshibkina 1983 = Ошибкина С. В. Мезолит бассейна Сухоны и Восточного Прионежья. Москва: Наука.

Oshibkina S.V. 1989. The Material Culture of the Veretye-type Sites in the Region to the East of Onega. – Ed. C. Bonsall. Clive The Mesolithic in Europe. Papers Presented at the Third International Symposium, Edinburgh, 1985. Edinburgh: John Donald, 402–413.

Oshibkina S.V. 1994. Mesolithic graves to east of the Onega Lake. – Archaeological News, 3. St.-Petersburg: Institute of the History of Material Culture, 48–57.

Oshibkina 1997 = Ошибкина С.В. Веретье 1. Поселение эпохи мезолита на Севере Восточной Европы. Москва: Наука.

Oshibkina 2004 = Ошибкина С.В. Мезолитическая культура веретье. Хронология и периодизация. – Российская археология, 1, 100–110.

Oshibkina 2006 = Ошибкина С.В. Мезолит Восточного Прионежья. Культура Веретье. Москва : Институт археологии Российской академии наук.

Oshibkina S. 2017. Funeral rituals of the population of the Eastern Lake Onega region (based on materials from Popovo and Peschanitsa cemeteries). – Mesolithic burials – Rites, Simbols and social organisation of early postglacial communities. Eds. J. M. Grünberg, B. Gramsch, L. Larsson, Y. Orschiedt & H. Meller. International Conference Halle (Saale), Germany, 18th-21st September 2013. Tagungen des Landesmuseums für Vorgeschichte Halle 13/I, Halle (Saale): Germany Verlag. Beier-Beran, 793–808.

Paavel, K., Kimber, A., Rannamäe, E., Kriiska, A. 2016. Investigations at Sõjamäe and Soodevahe Cup-marked Boulders and Late Neolithic. − Iron Age Settlement Site at the South-eastern Border of Tallinn. Archeological Fieldwork in Estonia 2015, 47−58.

Raushenbah 1960 = Раушенбах В. М. Фатьяновское погребение на неолитической стоянке Николо-Перевоз. Археологический сборнпк. Труды Государственного исторического музея, 37. Ed. Левашева В. П. Москва: , 28—37.

Saag, L., Varul, L., Lyn Scheib, C., Stenderup, J., Allentoft, M.E., Saag, L., Pagani, L., Reidla, M., Tambets, K., Metspalu, E., Kriiska, A., Willerslev, E., Kivisild, T., Metspalu M. 2017. Extensive Farming in Estonia Started through a Sex-biased Migration from the Steppe. Current Biology 27 (14), 2187−2193.

Sidorov & Engovatova 1992 = Сидоров В.В., Энговатова А.В. Ханевский могильник и древнейшие памятники фатьяновской культуры. – Археологические памятники Волго-Клязьминского междуречья, 7. Иваново, 28–37.

Tallgren, A.M. 1922. Zur Archäologie Eestis I. Vom Anfang der Besiedelung bis etwa 500 n. Chr. Acta et Commentationes Universitatis Tartuensis (Dorpatensis) B III: 6, Dorpat.

Tretyakov 1966 = Третьяков П.Н. Финно-угры, балты и славяне на Днепре и Волге. Москва–Ленинград: Наука.

Utkin, Kostyleva 2001 = Уткин А.В., Костылева Е.Л. Погребения на стоянке Караваиха. − Российская археология, 3, 55−66.

Uvarov 1881 = Уваров, А. С. Археологія Россіи. Каменный періодъ, I, II. Москва: Синодальная Типографія.

Varul, L., Galeev, R.M., Malytina, A.A., Tõrv, M., Vasilyev, S.V., Lõugas, L., Kriiska, A. 2019. Complex mortuary treatment of a Corded Ware Culture individual from the Eastern Baltic: A case study of a secondary deposit in Sope, Estonia. Journal of Archaeological Science: Reports 24, 463−472.

Volkova 2010 = Волкова Е. В. Новинковские могильники фатьяновской культуры. Москва: Институт археологии Российской академии наук.

Zaretskaya & Kostyleva 2011 = Зарецкая Н.Е., Костылева Е.Л. Новые данные по абсолютной хронологии льяловской культуры. – И.Н. Черных (ed.) Тверской археологический сборник, 8 (1). Тверь: ООО «Издательство «Триада», 175–183.

Zhilin et al. 2002 = Жилин М. Г., Костылева Е. Л,, Уткин А. В., Энговатова А. В. Мезолитические и неолитические культуры Верхнего Поволжья. По материалам стоянки Ивановское VII. Москва: Наука.

Zhilin M.G. 2003. Early Mesolithic communications networks in the East European forest zone – Eds. L. Larsson, H. Kindgren, K. Knutsson, D. Loeffler & A. Akerlund. Mesolithic on the Move: Papers Presented at the Sixth International Conference on the Mesolithic in Europe, Stockholm. Oxford: Oxbow books, 688–693.

Zhkov 1925 = Жуков Б.С. Неолитическая стоянка близ села Льялова Московского уезда. – Труды Антропологического института. 1. Приложение к Русскому антропологическому журналу, XIV (1–2). Москва, 37–78.

#### Supplementary Figures

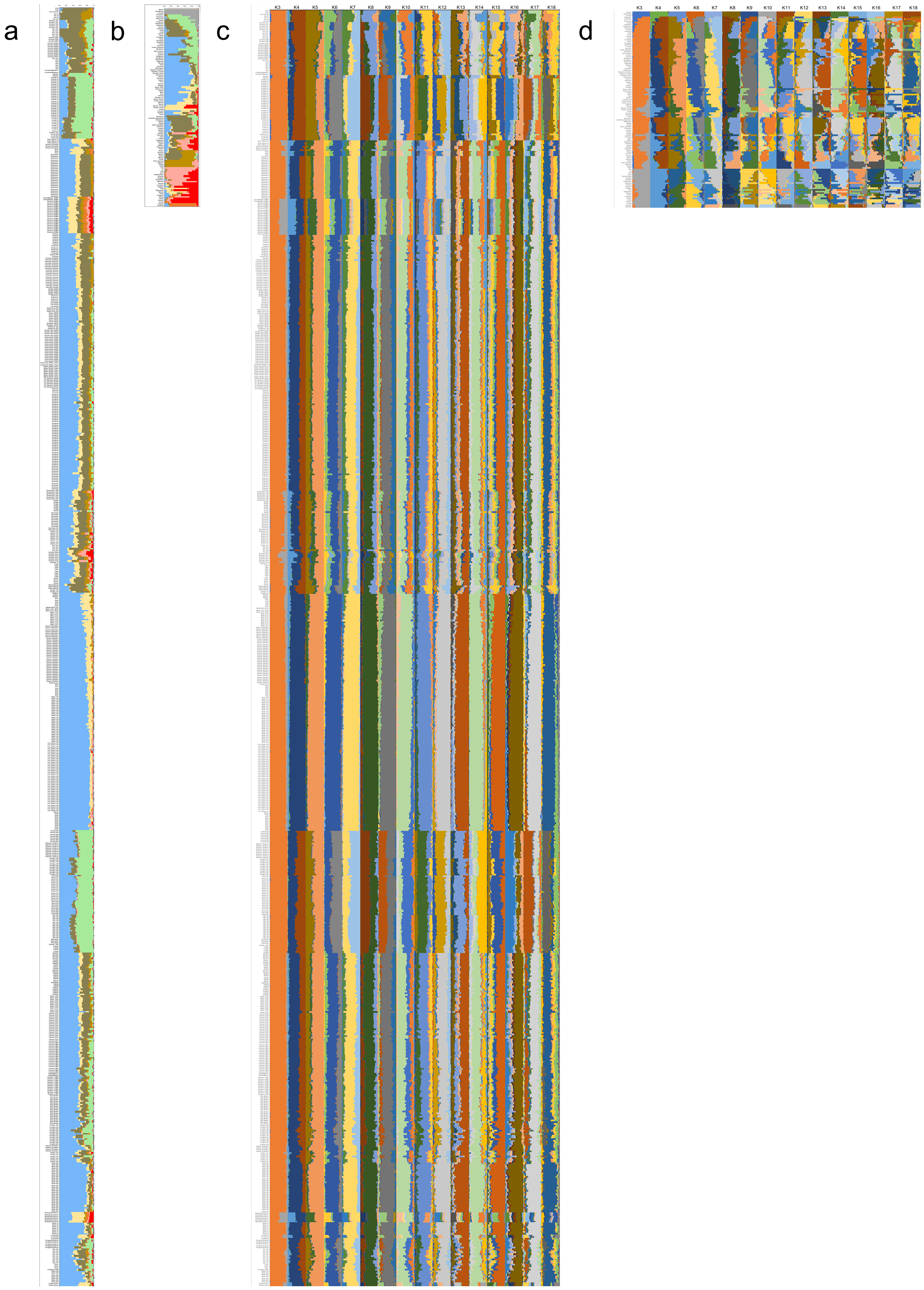


**Supplementary Figure 1. ADMIXTURE analysis results with ancient data projected onto the modern genetic structure. A** ancient individuals at K9, **B** modern population averages at K9, **C** ancient individuals at K3 to K18, **D** modern population averages at K3 to K18.

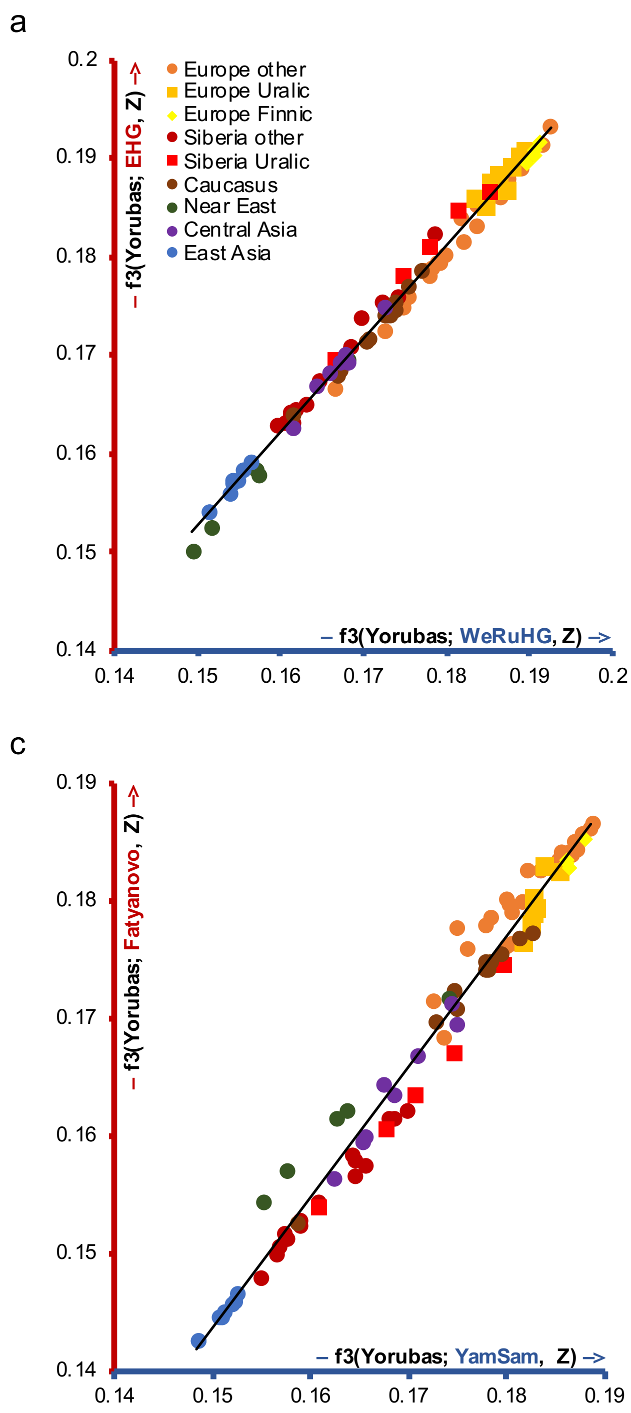


**Supplementary Figure 2. Outgroup f3 statistics' results of comparisons with modern populations.** Outgroup f3 statistics' values of form f3(Yorubas; study population, modern) plotted against each other for two study populations (blue and red axis): **A** Western Russian hunter-gatherers (WeRuHG) and Eastern hunter-gatherers (EHG), **B** WeRuHG and Fatyanovo, **C** Yamnaya Samara (YamSam) and Fatyanovo, **D** Central Corded Ware culture (CeCWC) and Fatyanovo.

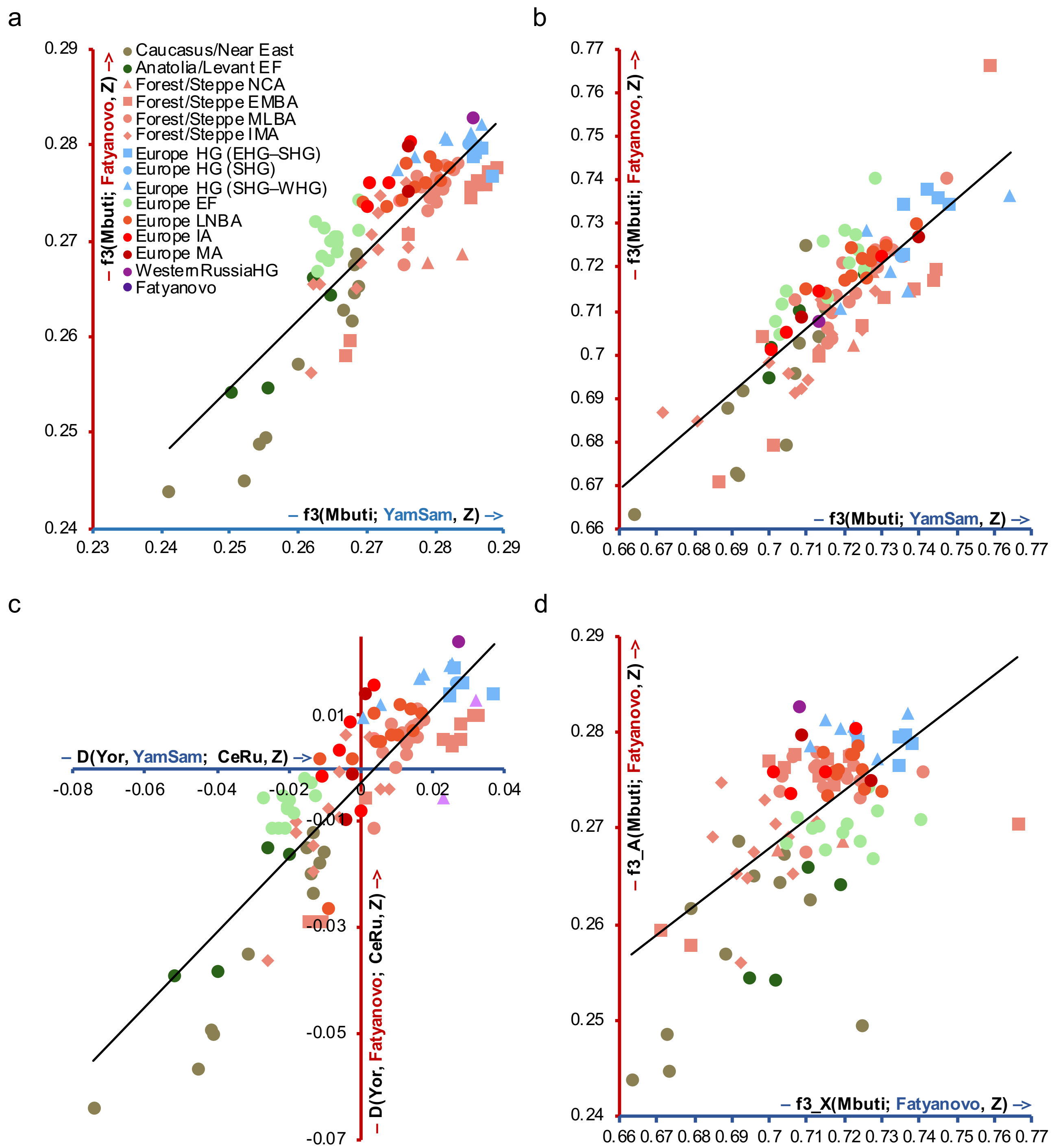


**Supplementary Figure 3. Outgroup f3 and D statistics' results of comparisons with ancient populations.** Outgroup f3 statistics' values of form f3(Mbuti; Yamnaya Samaya/Fatyanovo, ancient) using **A** autosomal, **B** X chromosome 1240k capture SNPs, **C** D statistics' values of form D(Yorubas, Yamnaya Samaya/Fatyanovo; Central Russians, ancient), **D** outgroup f3 statistics' values of form f3(Mbuti; chr X/autosomal SNPs, ancient) of Fatyanovo using 1240k capture SNPs. EF – early farmers; EMBA – Early/Middle Bronze Age; MLBA – Middle/Late Bronze Age; IMA – Iron/Middle Ages; HG – hunter-gatherers; LNBA – Late Neolithic/Bronze Age; IA – Iron Age; MA – Middle Ages.

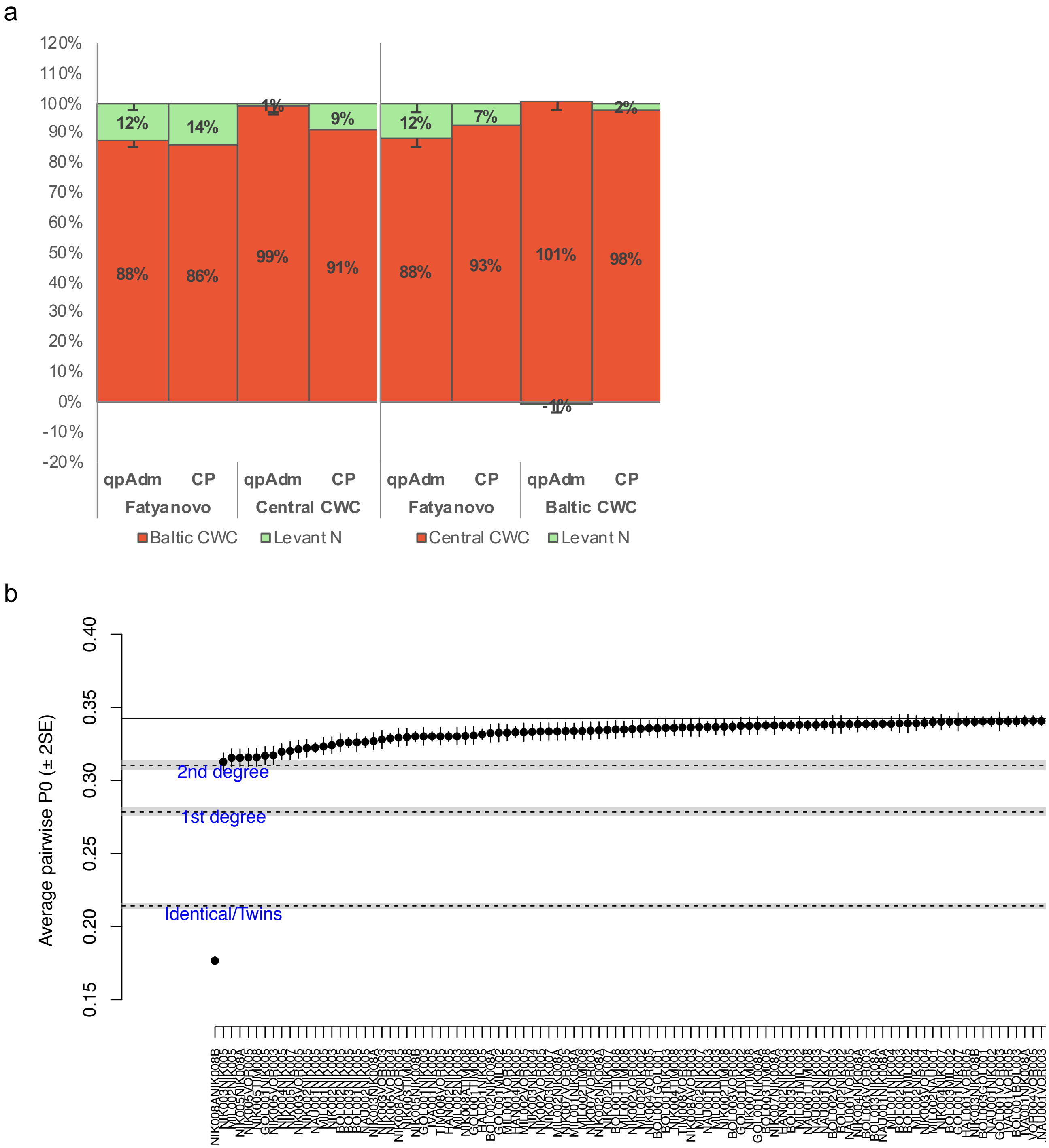


**Supplementary Figure 4. qpAdm, ChromoPainter/NNLS and kinship test results. A** qpAdm and ChromoPainter/NNLS (CP) models with Baltic Corded Ware culture (CWC) or Central CWC and Levant Neolithic (Levant N) as sources, **B** Fatyanovo kinship test results, 100 most similar pairs out of 253 shown.

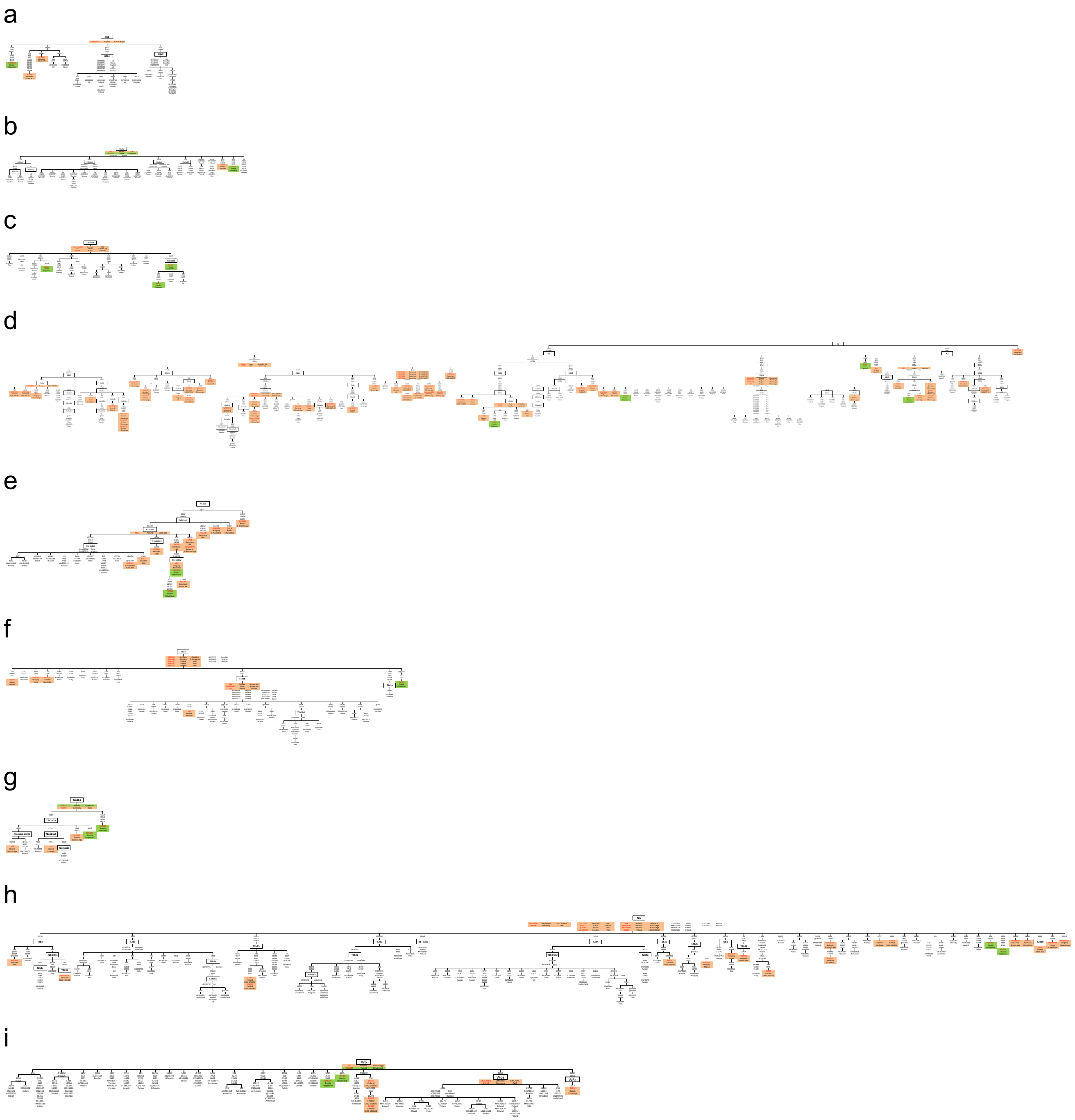

**Supplementary Figure 5. Partial mitochondrial DNA phylogenetic trees with modern and ancient individuals added. A** H5b, **B** H6a1a, **C** J1c1b1a, **D** K, **E** N1a1a1a, **F** T1a1, **G** T2a1b1, **H** T2b, **I** W6. Numbers on branches indicate substituted nucleotide positions consistent with Reconstructed Sapiens Reference Sequence (RSRS); only transversions are further specified with capital letters; @ refers to a reversion (back-mutations); .XC, .1C and .2C are insertions where X stands for unknown number. Modern data from https://www.ncbi.nlm.nih.gov/genbank/, https://www.internationalgenome.org/data; ancient from Allentoft et al. 2015, Fernandes et al. 2018, Furtwängler et al. 2020, Gamba et al. 2014, Haak et al. 2015, Juras et al. 2017, Knipper et al. 2017, Krzewinska et al. 2018, Linderholm et al. 2020, Malmström et al. 2019, Mittnik et al. 2018, Modi et al. 2019, Saag et al 2019, Valdiosera et al 2018, Översti et al 2019.
